# Epigenetic Control of Adaptive or Homeostatic Splicing of Alternative Exons and *Prolactin* Gene Expression During Interval-Training Activities of Pituitary Cells

**DOI:** 10.1101/2024.02.23.581772

**Authors:** Ling Liu, Hai Nguyen, Urmi Das, Samuel Ogunsola, Jiankun Yu, Lei Lei, Matthew Kung, Shervin Pejhan, Mojgan Rastegar, Jiuyong Xie

## Abstract

Interval-training activities induce adaptive cellular changes without altering their fundamental identity, but the precise underlying molecular mechanisms are not fully understood. In this study, we demonstrate that interval-training depolarization (ITD) of pituitary cells triggers distinct adaptive or homeostatic splicing responses of alternative exons. This occurs while preserving the steady-state expression of the *Prolactin* and other hormone genes. The nature of these splicing responses depends on the exon’s DNA methylation status, the methyl-C-binding protein MeCP2 and its associated CA-rich motif-binding hnRNP L. Interestingly, the steady expression of the *Prolactin* gene is also reliant on MeCP2, whose disruption during ITD leads to exacerbated overexpression and multi-exon aberrant splicing of the hormone gene transcripts, similar to the observed hyperprolactinemia or activity-dependent aberrant splicing in Rett Syndrome. Therefore, depending on how many times cells are stimulated, exons may exhibit different splicing responses to cell activities. During the ITD, epigenetic control is crucial for both adaptive or homeostatic splicing and the steady expression of the *Prolactin* hormone gene. Disruption in this regulation may have significant implications for the development of progressive diseases.

## Introduction

Interval-training activities confer beneficial effects on cardiac, muscular and endocrine functions that cannot be attained through a single bout of activity ^1–4^. Despite this, the underlying molecular and cellular mechanisms for the adaptive as well as homeostatic responses of different genes remain to be fully elucidated. The regulation of gene expression is critical in facilitating adjustments essential for functions such as hormone synthesis ^2,5,6^. However, the regulation of alternative pre-mRNA splicing^7^, a fundamental mechanism driving the diversification of metazoan transcriptomes and proteomes to support intricate cellular functions ^8,9^, remains enigmatic in the adaptation process.

We have explored how cellular activities, particularly through membrane depolarization, regulate alternative splicing in pituitary cells^10–13^, where we predicted that cells adapt their splicing patterns after repeated stimulation compared to initial treatments^12^. This regulation has been shown to be crucial for the alternative splicing of various genes in response to chronic changes in membrane potentials, significantly affecting neuronal electrical homeostasis or synaptic formation^14–16^. The regulatory mechanisms involve Ca^++^/calmodulin-dependent protein kinase IV (CaMKIV) and downstream splicing factors such as hnRNP L/LL^10–12,17^, Sam68^15^,or Nova-2^14^, depending on the target exon. Additionally, histone modifications and the methyl-DNA-binding protein MeCP2 play a key role in activity-dependent regulation, suggesting epigenetic influences^18–20^. However, the role of DNA methylation in this process remains unclear.

DNA methylation is pivotal in adaptation^1,21^, and generally correlates with exon inclusion in the genome/transcriptome ^21^, though not in all cases^22^. The correlation aligns with the regulatory effects of MeCP2 and the methyl-free DNA-binding CTCF on splicing^23–25^. MeCP2, which binds to specific nucleotide sequences mCAC and mCG^26^, influences gene transcription and splicing ^25–27^. Notably, MECP2 mutations are identified in up to 96% of typical Rett syndrome cases^28–30^, a severe neurodevelopmental disorder with autistic features and often exacerbated by abnormal brain activities like epilepsy^31^. In Mecp2-null mouse models of Rett syndrome, long genes’ expression and synaptic exon splicing in the hippocampus, particularly after calcium signal-activating kainic acid treatment, are significantly impacted^19,32^. Interestingly, both DNA methyltransferase DNMT3a and MeCP2 are regulated by calcium signaling: DNMT3a is recruited by CaMKIV-regulated CREMα in T lymphocytes^33,34^, and MeCP2 is phosphorylated by CaMKII in hippocampal neurons^35^. MeCP2 likely modulates alternative splicing through its interaction with methylated DNA and splicing factors like YB-1^25,36^. Together, DNA methylation/MeCP2 dysfunction likely play important roles in the development of neurological diseases as many other epigenetic changes^37^. However, bioinformatics analyses suggest minimal global effects of DNMTs and MeCP2 on alternative splicing^38^. These inconsistencies highlight the need for further research using comprehensive methylation and splicing analyses, including direct methylation of exon DNA in splicing assays, to clarify the effects of DNA methylation and MeCP2 on splicing.

Here we show the distinct effects of interval-training depolarization (ITD) compared to a single round of treatment on the adaptive splicing of exons in the prolactin and growth hormone-producing pituitary cells^39^. We identified a critical role of epigenetic control in both adaptive or homeostatic splicing and *Prolactin* gene expression, a dual function in adaptation while preserving cell identity.

## Results

### 1. Adaptive splicing induced by ITD of GH_3_ pituitary cells and its disruption by the DNA methylation inhibitor 5-azacytidine

The depolarization effect on splicing is reversible by washing off/adding back depolarizing concentrations of KCl (50mM)^40^. We thus mimicked interval-training activities of cells by treating GH_3_ pituitary cells with interval-training depolarization (ITD), in comparison to the single round of treatment established in these cells in our previous studies^11,12^. The cells were treated once or six times with depolarizing concentrations of KCl for 6h, followed by 18h wash-off intervals (Fig. 1A, see also S_Fig. 1A-B for pre-tests of the STREX exon^10^). Our treatment did not alter the growth curve of the cells (S_Fig. 1C), and showed strong homeostatic expression for the majority of exons (99%, S_Fig. 1D) and the signature *Prolactin* and *Growth hormone* genes of the pituitary cells ^41^ (see below).

**Figure 1.**
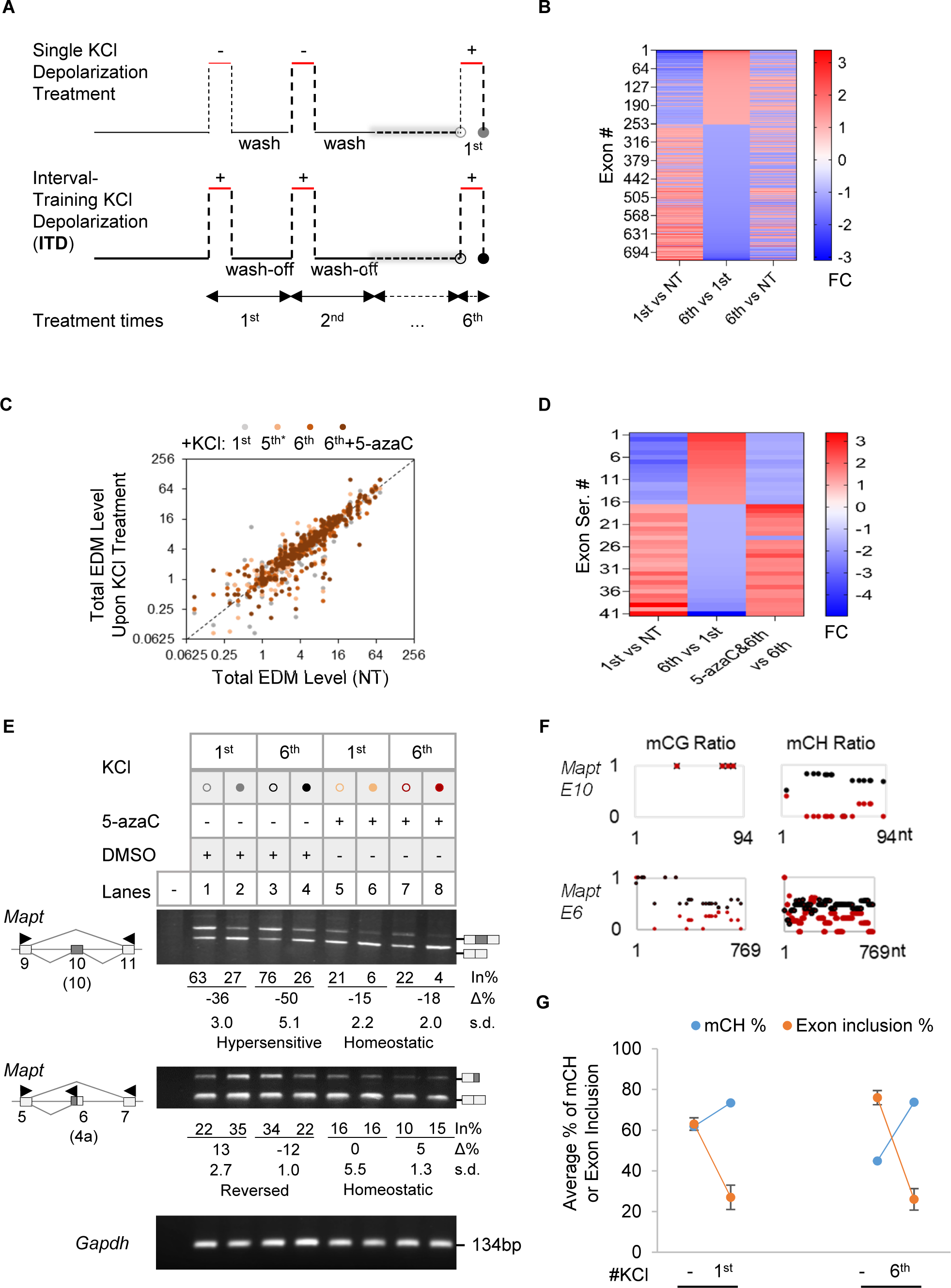
Adaptive splicing of a group of exons upon ITD of GH_3_ pituitary cells and effect of the DNA methylation inhibitor 5-azaCytidine. **A.** Scheme of the single or ITD KCl (50mM) treatments of GH_3_ pituitary cells. **B.** Heatmap of the fold changes (FC) of 720 exons between the 6^th^ and 1^st^ KCl-treated cells by DEXSeq analysis (≥ 1.1-folds, p < 0.01, average base mean >20, exon base mean >20). Additional exons were detected by MATS analysis. **C**. EDM levels upon the 1^st^ or ITD KCl treatments versus their levels in NT samples by BSMAP analysis of whole genome bisulfite sequencing (WGBS) data (n = 346 exons with measurable mC in all samples). Grey, yellow, orange or red dot: 1^st^, 5^th^, 6^th^ and 6^th^ KCl plus 5-azaC treatment, respectively. *: 18h after wash-off of the 5^th^ KCl media, before the 6^th^ KCl addition. NT, not treated. **D.** Heatmap of the usage fold changes of 41 top adaptive exons (FC > 1.5) that are prevented by 5-azaC (FC > 1.5). **E.** Representative agarose gels of RT-PCR products (Left) of 5-azaC effects on the adaptive splicing of exons in response to single round of depolarization or ITD treatments (n ≥ 3). Arrowheads: PCR primers. The exon numbers are based on reference transcripts in the UCSC Genome Rat Jul. 2014 (RGSC 6.0/rn6) Assembly: Mapt exon 6, M84156, equivalent to human MAPT exon 4a (NM_001123066.3, GRCh38/hg38); Mapt exon 10, M84156, equivalent to human MAPT exon 10 (NM_001123066.3, GRCh38/hg38). In all the exams, the delta scores showed significant differences, except for those in Mapt exon 6(4a) during either the 1st or 6th KCl treatment with 5-azaC. **F.** The corresponding mC ratios of the exons in **E** upon the 6^th^ KCl treatment with (red) or without (black) 5-azaC. **G.** An example of EDM changes (*Mapt* exon 10, in 100% scale) by the 1^st^ and 6^th^ depolarization treatment, alongside its exon inclusion levels.

A subset of exons (1,878 exons of 1,204 genes) displayed substantial changes (Fig. 1B). Around 81% of them were alternatively spliced exons and the remainder were produced from alternative transcription start or polyadenylation sites. These genes mainly clustered for functions at the synapse or for RNA recognition (S_Fig. 1E). Their alternative exons exhibited three primary response patterns upon ITD: homeostatic, desensitized, or hypersensitive compared to their responses to the single (1^st^) KCl treatment (S_Fig. 1F-H). Validation by semi-quantitative reverse transcription-polymerase chain reaction (RT-PCR) confirmed adaptive splicing in 78.3% of exons (n = 23) examined by their altered net percent changes of exon inclusion upon ITD over the 1^st^ treatment (S_Fig. 1G-H, desensitized or hypersensitive, *p* < 0.05 by Δ%). Therefore, ITD induces adaptive splicing of a group of exons in genes associated with key cellular functions amidst the homeostatic response of most exons.

In subsequent tests of several pathway inhibitors including the known calcium signaling inhibitor nifedipine in the depolarization-regulated splicing^13^ (S_Fig. 2), 5-azacytidine (5-azaC), an inhibitor of DNA methyl-transferases (DNMT) ^42,43^, significantly disrupted splice site usage^44^ and the adaptive splicing of endogenous exons (Fig. 1C-F, and S_Fig. 3). The adaptive synaptic exons (S_Fig. 1E) together with epigenetic effects in neurological diseases^37^, also made the 5-azaC effect an interesting direction for further investigation.

**Figure 2.**
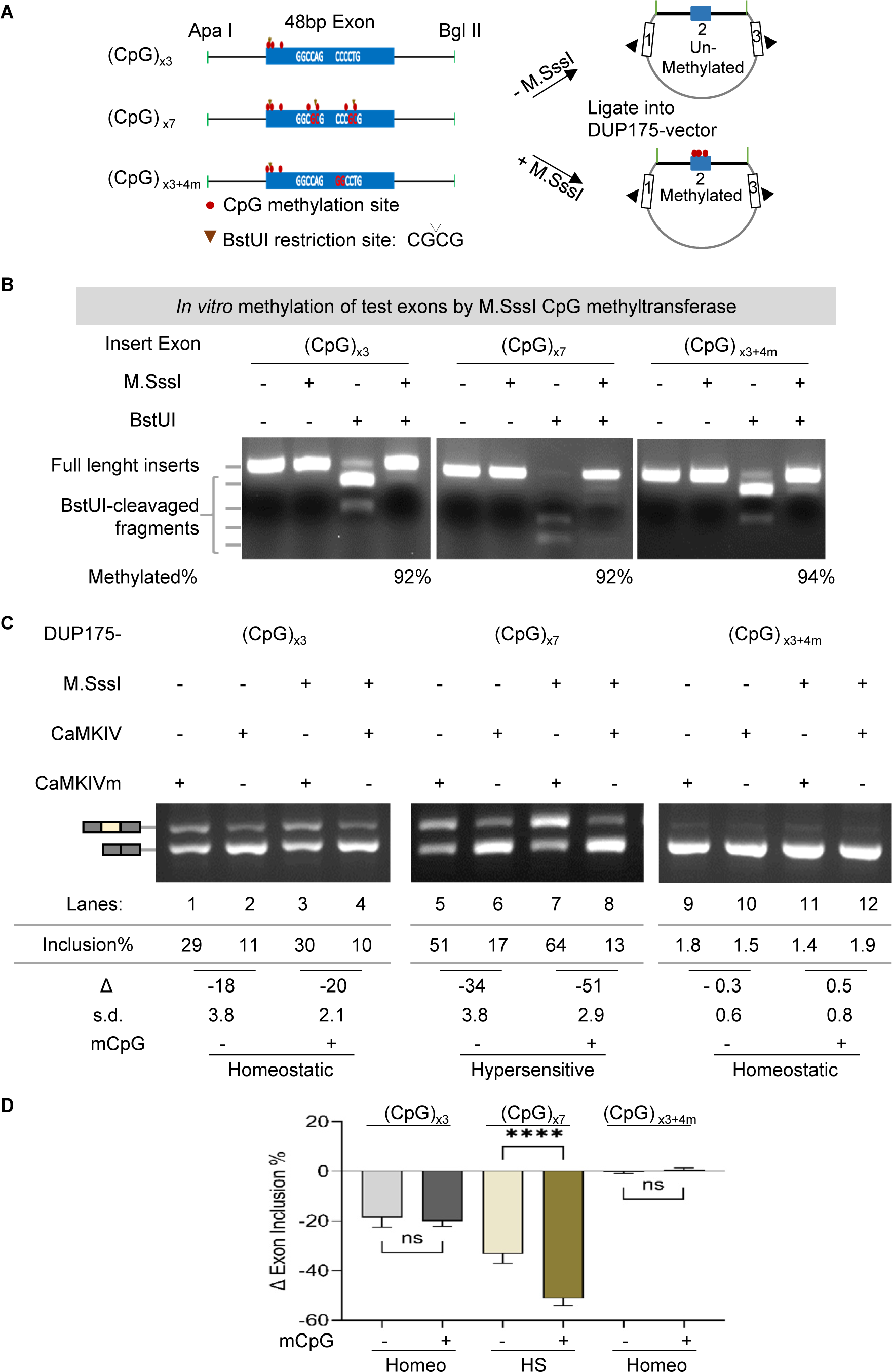
Effect of the EDM status on the adaptive or homeostatic splicing in response to CaMKIV. **A.** Diagram of the *in vitro* EDM minigene splicing reporter assay. The synthesized reporter exons are based on the adaptive exon 14 (48nt, ENSMUST00000106233.1, GRCm38/mm10) of the mouse *Baiap2* gene (see below). Its partial flanking introns here do not harbor CpG sites. The vector backbone is the DUP175, derived from the constitutive beta-globin exons. Arrowheads: PCR primers. **B.** The *in vitro* methylation efficiency of the reporter exons [(CpG)_x3_, (CpG)_x7_ or (CpG)_x3+4m_] by M. SssI CpG methyltransferase, verified by using the CpG methylation-sensitive BstUI (restriction site: CG↓CG). The percentage of the full-length insert (minus the uncleaved full-length, unmethylated insert in the preceding lane) out of all fragments in each lane was taken as the methylation efficiency. **C.** Agarose gels of semi-quantitative RT-PCR products of the splicing reporters transiently expressed in HEK293T cells with CaMKIV or its mutant (CaMKIVm). **D.** Bar graph of net percent changes of the reporter exons in **C** by CaMKIV (n ≥ 3). Homeo: homeostatic; HS: hypersensitive; ns: not significant; ****: *p* < 0.0001.

**Figure 3.**
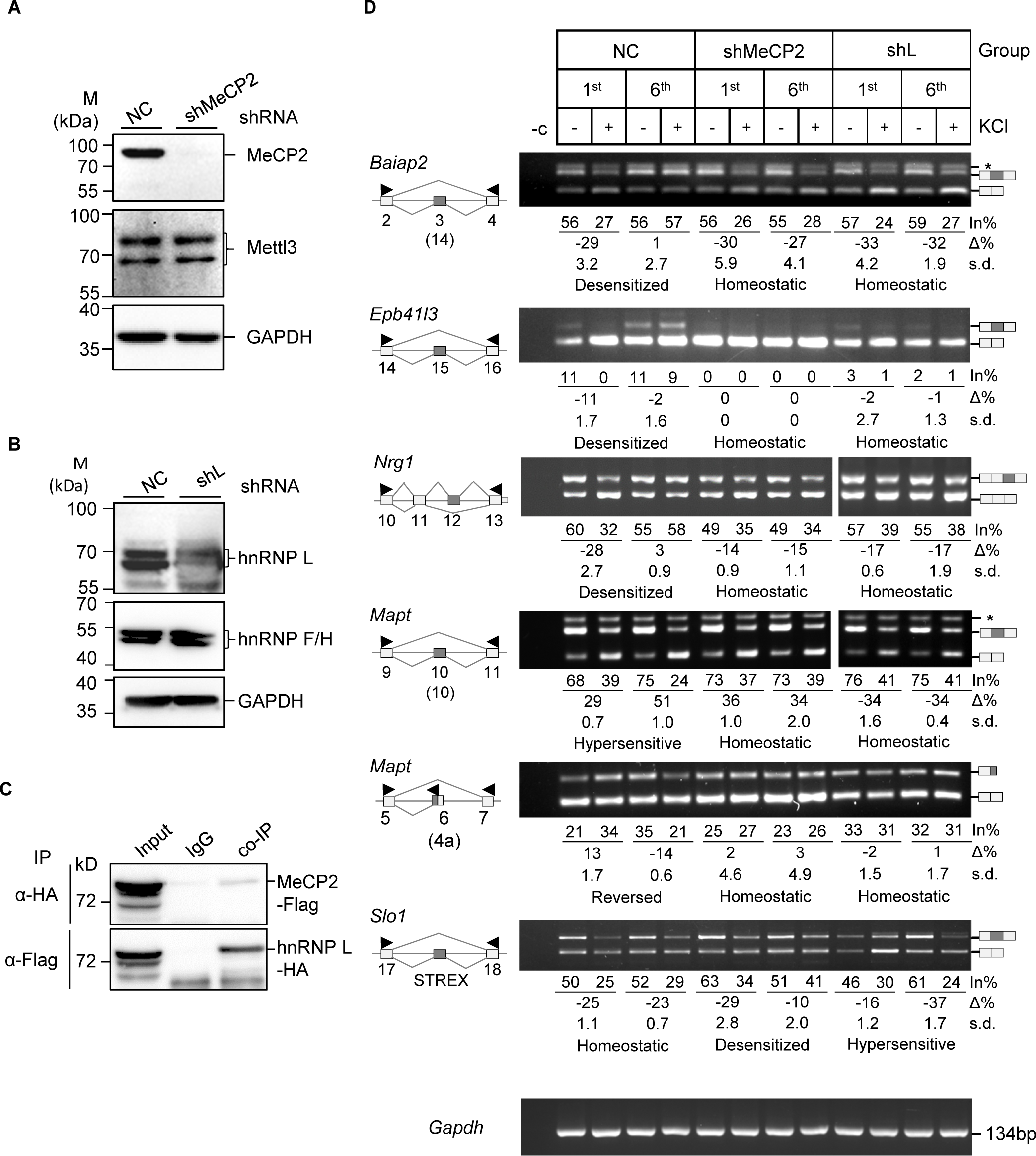
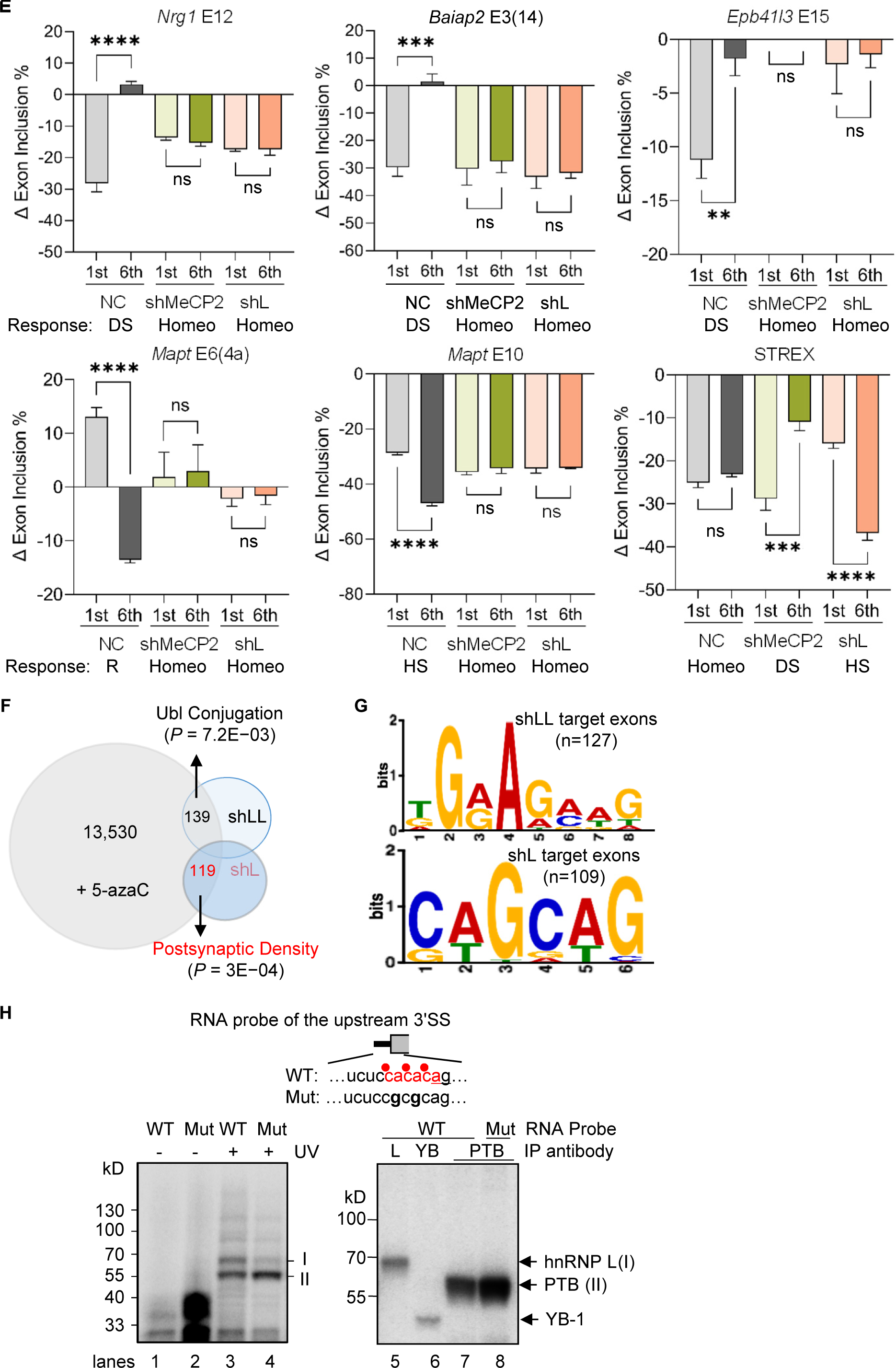
Effect of knocking down MeCP2 or its partner hnRNP L on adaptive or homeostatic splicing. **A-B.** Western blot analyses of stable lentiviral shRNA-expressing GH_3_ cell lines showing specific knockdown of MeCP2 (**A**) or hnRNP L (**B**) proteins compared to the non-targeting control shRNA (NC). Mettl3, hnRNP F/H and GAPDH are shRNA-negative and protein loading controls. **C.** Reciprocal co-immunoprecipitation assay of MeCP2-Flag and hnRNP L-HA co-expressed in HEK293T cells. IgG: rabbit IgG, negative control. **D.** Representative agarose gels of RT-PCR products of the major patterns of MeCP2 or hnRNP L knockdown effects on the adaptive or homeostatic response of exons that are also disrupted by 5-azaC (n ≥ 3). Arrowheads: PCR primers. **E.** Bar graphs of net percent changes of adaptive or homeostatic exons upon single (1^st^) or ITD (6^th^) KCl treatment, with/without MeCP2 or hnRNP L knockdown (n ≥ 3). Homeo: homeostatic; DS: desensitized; HS: hypersensitized; R: reversed. ns: not significant, **: *p* < 0.01; ***: *p* < 0.001; ****: *p* < 0.0001. **F.** Diagram of differentially spliced exons in the GH_3_ transcriptome upon 5-azaC treatment or lentiviral knockdown of hnRNP L protein by shL, with the L-like (LL) targets in comparison. **G.** Consensus motifs of shLL- or shL-changed exons that were also affected by 5-azaC, identified by MEME analysis. **H**. UV crosslinking of wild type (WT) CA repeat or CG mutant (Mut) RNA probes of the *Mapt* exon E6 upstream 3’ splice site in HeLa nuclear extracts. Upper: probe diagram. Red dots: corresponding DNA mCpAs, of which methylation levels were reduced by 5-azaC. Underlined: 3’ AG. Lower: phosphorimages of proteins crosslinked to the probes and resolved in SDS-PAGE gels. Immunoprecipitating antibody is against hnRNP L (L), YB-1 (YB) or PTB (PTBP1). A sixth of the crosslinking mix for immunoprecipitation was loaded in lanes 3 and 4.

Specifically, prior addition of 5-azaC to the cells before the 6^th^ KCl treatment led to global alterations in exonic DNA methylation (EDM) and exon usage, which inversely correlated with each other overall (S_Fig. 3A-B). Consistent with these effects, KCl depolarization itself indeed also caused EDM changes in across 346 exons detected along the course of the treatments compared to that of the non-treated (Fig. 1C). Importantly, 41 of the top 52 adaptive exons (∼79%) were disrupted by at least 1.5 folds when treated with 5-azaC (Fig. 1D). RT-PCR analysis confirmed that the adaptive splicing of these exons was mostly abolished to homeostatic responses by 5-azaC (Fig. 1E and S_Fig. 3C), and two of their response patterns reversed (Dlg and Phldb1 exons). More interestingly, we also noticed that even some homeostatically responsive exons became adaptive upon the 5-azaC treatment (e.g. STREX in S_Fig. 1G and Mapt E7a in S_Fig. 3C). The 5-azaC effect on the patterns of splicing responses to ITD was accompanied by EDM changes (Fig. 1F-G, and S_Fig. 3D). Together, these data suggest that EDM probably play a role in the ITD-induced patterns of adaptive or homeostatic splicing in an exon-specific manner.

### 2. EDM level-dependent adaptive or homeostatic splicing of *in vitro* methylated exons

To directly explore the influence of EDM on adaptive or homeostatic splicing, we created synthetic reporter exons harboring either 3 or 7 copies of CpG dinucleotides that can be specifically methylated *in vitro* (Fig. 2A, from exon 14 of the synaptic m*Baiap2* gene^45^). The methylation efficiency reached at least 90% (Fig. 2B) by the M. SssI CpG methyltransferase ^46^. The reporters with various EDM levels were tested with co-expressed CaMKIV, a key mediator of depolarization-induced splicing ^10,12,13,15,40^.

Upon co-transfection into HEK293T cells with the constitutively active Flag-CaMKIV-dCT or its kinase-dead mutant^12,13,17^, we observed that the (CpG)_×3_ exon was repressed by CaMKIV by approximately 20% reduction, regardless of its methylation status, consistent with a homeostatic response (Fig. 2C, lanes 1-4). However, increasing the CpG count to 7, resembling the ITD-induced EDM changes (Fig. 1C, F-G), augmented the CaMKIV repression from 34% to 51% reduction upon hypermethylation, consistent with a hypersensitive response (lanes 5-8). In contrast, mutating the additional 4 copies of CpG dinucleotides eliminated this enhancement effect, reversing it back to homeostatic response (lanes 9-12). This indicates that the EDM level determines the exon’s homeostatic (lanes 1-4, 9-12) or adaptive (lanes 5-8) response to the same stimulation by the co-expressed CaMKIV, and the EDM change is both necessary and sufficient for an altered/adaptive splicing response to CaMKIV. Taken together with the ITD-induction of EDM changes and their correlation with the splicing changes (Fig. 1C, G and S_Fig. 3A-B, D), these results support an essential role of EDM level and its changes in ITD/CaMKIV-induced adaptive or homeostatic splicing.

### 3. Essential role of MeCP2 and its partner hnRNP L in ITD-induced adaptive or homeostatic splicing of a group of exons

Based on the EDM analysis, the roles of the mC-binding MeCP2 and CaMKIV-target splicing factor hnRNP L in activity-dependent splicing^10,19^, and synaptic gene exons changed by both 5-azaC and hnRNP L^47^ (S_Table I), it’s likely that MeCP2 and hnRNP L are involved in the ITD-induced splicing patterns as well. We thus established GH_3_ cell colonies stably knocked down of MeCP2 or hnRNP L using lentiviral shRNAs (Fig. 3A-B). Interestingly, these two proteins interact in reciprocal co-immunoprecipitation assays (Fig. 3C). Upon knockdown of either MeCP2 or hnRNP L, all five adaptive exons of the synaptic genes tested lost their adaptive splicing patterns to homeostatic responses to ITD (Fig. 3D-E), in a similar way as the same *Epb41l3*, *Nrg1* and *Mapt* exons by 5-azaC in S_Fig. 3C and Fig. 1E. The homeostatic STREX exon (see also S_Fig. 1G), in contrast exhibited desensitized or hypersensitive responses to ITD upon knockdown of the two proteins, respectively.

In further support of hnRNP L’s role in the control of additional exons besides the reported STREX^10,17,47^, we examined the sequences of the 119 exons regulated by both hnRNP L and 5-azaC (Fig. 3F and S_Table I). They share specifically an hnRNP L-preferred CA-rich consensus motif in or nearby the exons (Fig. 3G). Moreover, UV-crosslinking-immunoprecipitation and CA-to-CG mutation assays of the *Mapt* exon 6 motif at the 3′ splice site supported hnRNP L direct binding to the site in a CA-dependent manner (Fig. 3H). In the WGBS analysis (Fig. 1C), the mC ratio around the 3′ splice site of the Mapt exon 6 (4a) changed from 0.75 to 0.54 by the 1^st^ and 0.66 to 1.0 by the 6^th^ KCl treatment, respectively, and then reduced to 0.29 by 5-azaC.

Together, these findings support that both MeCP2 and its associated hnRNP L are required for either adaptive or homeostatic splicing responses of the synaptic alternative exons upon ITD.

### 4. Essential role of MeCP2 for the homeostatic expression and proper splicing of the *Prolactin* hormone gene upon ITD of the pituitary cells or for the proper splicing of the adaptive exons of synaptic genes in the hippocampus of Rett syndrome patients

In addition to disrupting ITD-dependent adaptive or homeostatic splicing, we also found that treatment with 5-azaC exacerbated aberrant splicing of a number of hormone or hormone-related genes including the *Prolactin* gene upon ITD, accompanied by EDM changes (S_Figs. 4-6). MeCP2 knockdown slightly increased the *Prl* transcript level but more prominently, exacerbated aberrant splicing in an ITD-dependent way (Fig. 4A), causing the skipping of constitutive exons 2 and 3 coding for the conserved domain of the growth hormone-like superfamily^47,48^, or the inclusion of a cryptic 93nt exon found in hnRNP L-knockdown cells^47^. However, the growth hormone gene expression and splicing were not affected in these cells, indicating that the homeostatic transcript level and proper splicing of the pituitary signature gene *Prolactin* specifically requires sufficient MeCP2 in response to ITD. This finding also aligns closely with the *MECP2* mutation-aggravated aberrant splicing induced by chronic neuronal activities^19^, as well as with abnormal neuronal activities and synaptic plasticity associated with the progression of Rett syndrome ^31,49–52^. Moreover, the increased *Prl* mRNA transcripts upon MeCP2 knockdown may help explain the origin of the hyperprolactinemia in about 14% of the Rett syndrome patients^53,54^. Together our findings support that MeCP2 is critical for the maintenance of homeostatic level and the prevention of cryptic splicing for the integrity of the prolactin hormone transcript upon ITD in the pituitary cells.

**Figure 4.**
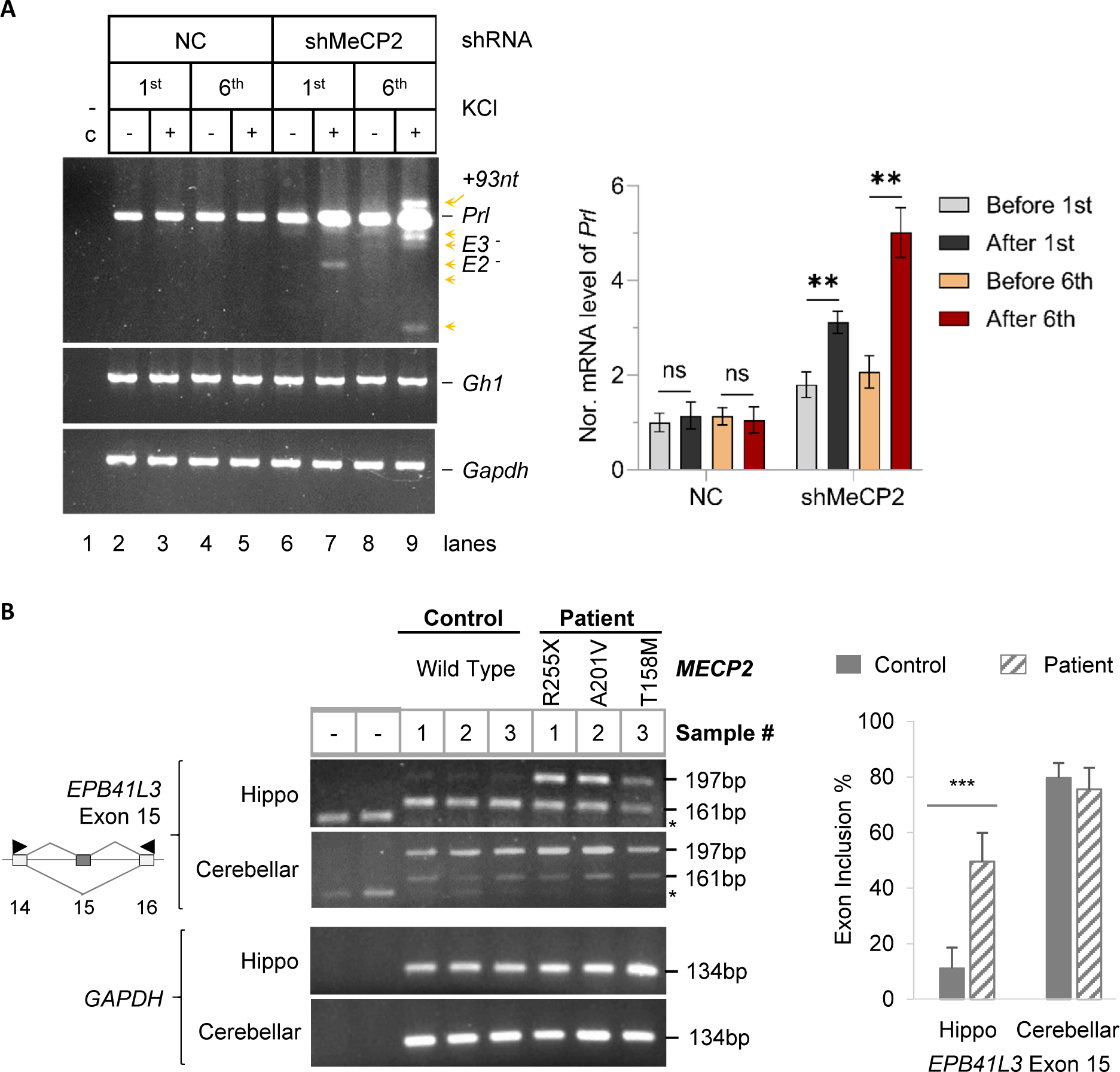
Exacerbated overexpression and aberrant splicing of the *Prolactin* gene in MeCP2-knockdown GH_3_ cells upon ITD and aberrant splicing of the adaptive exons of synaptic genes in Rett syndrome patients. **A.** RT-PCR products of the hormone *Prolactin* (*Prl*) gene upon MeCP2 knockdown and ITD treatment of GH_3_ pituitary cells. *Gh*: growth hormone gene as a negative control of the hormone genes; *Gapdh*: RNA loading control. Arrowheads: aberrant splice variants with the deduced cryptic skipped/included exons to the right. +93nt: inclusion of a 93nt cryptic exon in the *Prl* intron 4 as reported by Lei, et al., *MCB*′18. RNA samples are as in Figure 3A and D. The bar graph to the right shows the ITD-dependent increase of the index level of the *Prolactin* mRNA transcripts normalized to *Gapdh*. **B**. (Left) Agarose gel RT-PCR products of the adaptive *EPB41L3* synaptic gene/exon in the hippocampal tissues (Hippo) of Rett syndrome patients and healthy controls. Cerebellar tissues of the same patients are used as controls. *: Product from primers without cDNA. (Right) Bar graphs of the percentages of exon inclusion (mean ± s.d., n = 5 Rett syndrome patients, and 4 healthy controls). ns: not significant; **: *p* < 0.01; ***: *p* < 0.001.

To corroborate these *in vitro* findings from cultured cells, we evaluated the splicing of the ITD-induced adaptive exons in MeCP2-defective mouse or human samples. In *Mecp2*-null mice ^19,55^, the calcium signal-activator kainic acid (KA) generally exacerbated splicing changes (S_Fig. 7A), consistent with previous studies ^19^. Interestingly, these splicing alterations were also inversely correlated with EDM globally, including 8 of the top adaptive exons affected by 5-azaC (from Fig. 1D). Importantly, we identified several of the adaptive exons in Rett syndrome patients with *MECP2* mutations ^29,50,56^. These exons, particularly the synaptic *EPB41L3* exon 15, showed significant splicing changes in the hippocampi but not cerebella (Fig. 4B, and S_Fig. 7B-D), indicating hippocampus-specific aberrant splicing of the ITD-induced adaptive exons upon loss-of *MECP2* function.

Taken together, the epigenetic control is likely essential for both adaptive or homeostatic splicing of alternative exons and the homeostatic expression and proper splicing of the signature *Prolactin* hormone gene upon ITD of pituitary cells, with implications for the aberrant splicing of such exons in the progressive genetic disease Rett syndrome.

Traditionally, studies on gene expression and particularly alternative splicing have focused on the effects of single or continuous treatments. However, cells often experience repeated extracellular stimuli interrupted by periods of inactivity, as seen in neurons, muscle cells and hormone-producing pituitary cells in such activities as interval-training exercises or drug addiction ^1,2,5^. Our study extends the existing body of work by examining how cells adapt their alternative splicing pattern in response to interval-training depolarization (ITD) while maintaining their identity by homeostatic gene expression and splicing responses. The results here demonstrate clearly that exons may exhibit different responses to cell activities depending on how many times the cells are stimulated. The altered patterns of splicing responses would likely allow cells to fine-tune their adaption during such activities as exercises or stress in endocrine cells^2,3,5,6^, and to electrical firing activities in neurons ^11,12,14–16^. The dysregulation of the adaptive/homeostatic splicing upon epigenetic disruption (Fig. 4) likely contribute to the aberrant gene expression or splicing in the progression of neurological diseases. The diverse effects of the epigenetic control of different exons probably suggest an important role of the exon-dependent interplay between the epigenetic and splicing machineries (Fig. 5) in adaptive and homeostatic cell physiology and progressive diseases. The interplay is worthy of further study for the exon-dependent effect and underlying molecular mechanisms that may involve more epigenetic/splicing factors in the process (S_Fig. 8 and S_Tables II and III).

**Figure 5.**
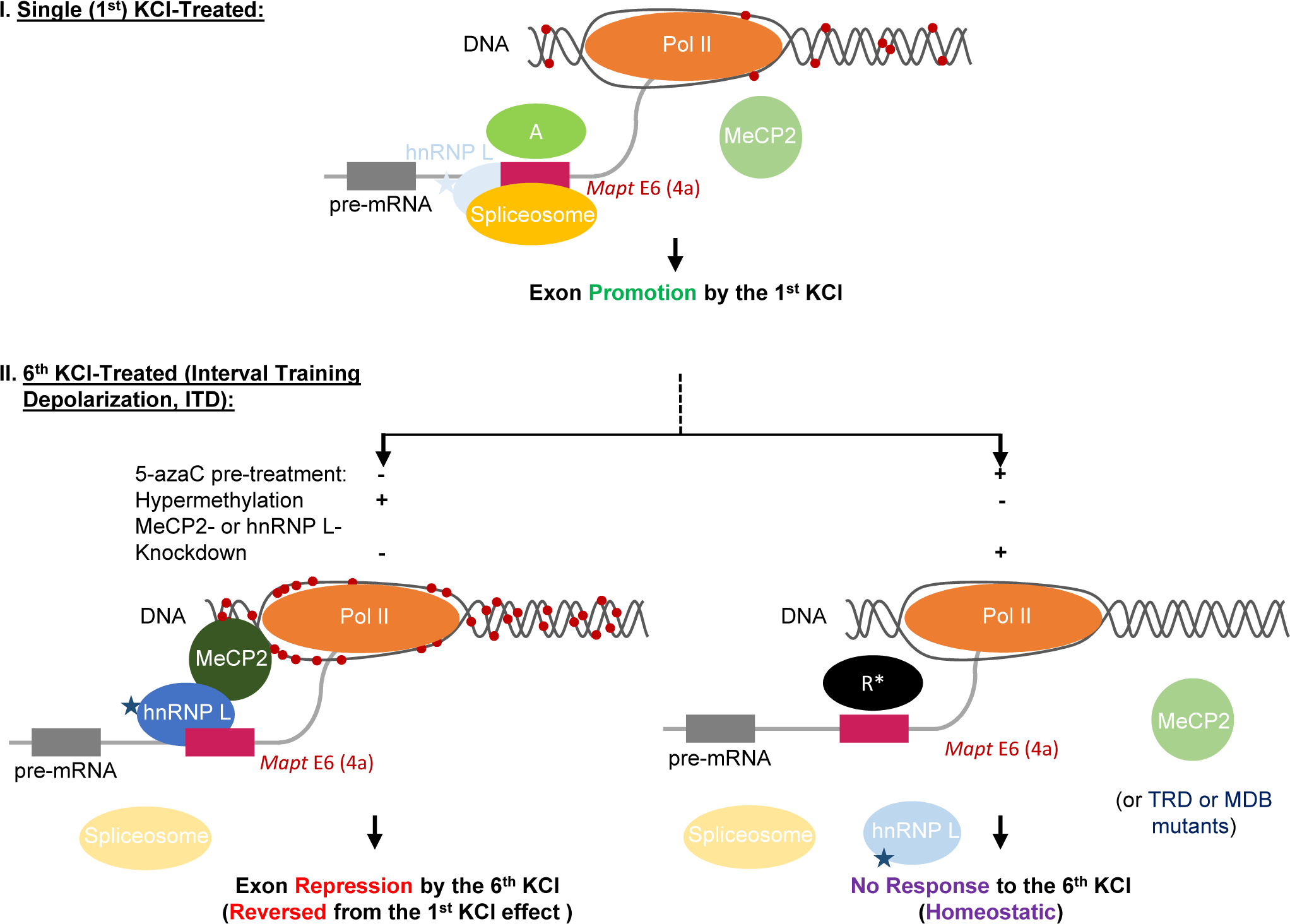
Summary of the epigenetic components in the control of adaptive or homeostatic splicing upon ITD. Shown are effects on the *Mapt* exon 6 (4a) as an example of the exon-dependent, diverse changes of methylation/splicing. With a single round of KCl treatment (**I**), the DNA methylation at the 3′ splice site is at a comparatively low level (54%); therefore, binding of MeCP2 and its associated hnRNP L to the DNA and pre-mRNA is inefficient, and insufficient to overcome the effect of splicing activators (A) in cells. The 6^th^ KCl-treatment (**II**, ITD), the DNA is hypermethylated (100%, Left panel) recruiting MeCP2 and its associated hnRNP L leading to exon repression (reversed splicing response from the 1^st^ KCl treatment). Right panel: with 5-azaC pre-treatment (Fig. 1E-F), the methylation is greatly reduced (29%) and the exon is inhibited by a depolarization non-responsive repressor (**R**) thereby homeostatic splicing response. *: In case of MeCP2- or hnRNP L-knockdown (Fig. 3D), the repressor R is likely replaced by a depolarization-non-responsive activator also causing homeostatic splicing though with higher basal level of the exon. Similar decoupling of the methyl-DNA/MeCP2 and associated hnRNP L is likely most effective when the TRD and MDB domains of *MECP2* are mutated (S_Fig. 7D). The mC changes and their disruption during the adaptive splicing are also consistent with the *in vitro* methylation/mutagenesis data of the reporter exon in Fig. 2. Star: p-Ser^513^ of hnRNP L by CaMKIV. Red dots on DNA: methyl-Cytosines. Lighter colors of the factors/spliceosomes represent their reduced effects.

## Materials and Methods

### Cell culture

Rat GH_3_ pituitary cells were cultured at 37 °C with 5% CO_2_ in Ham’s F10 nutrient mixture with 10% horse serum plus 5% fetal bovine serum (FBS), and human embryonic kidney (HEK) 293T cells in DMEM with 10% FBS. Penicillin-streptomycin-glutamine solution was added to all cultures except GH_3_ (without glutamine). HEK293T cells were dispersed by trypsin (0.05%, w/v) - EDTA (0.53 mM) solution during subculture.

### Interval-repeat KCl treatments and RNA/DNA extraction

For the ITD KCl treatment group, GH_3_ cultures were treated with KCl (50mM) for 6h, then washed and supplied with complete fresh medium, followed by incubation for 18h, completing the 1^st^ round (day) of the KCl treatment, which was repeated up to the 6^th^ time (6^th^ KCl). For the single KCl treatment group (1^st^ KCl), cells went through the same medium change process except that KCl (50mM) was added on the 6^th^ day. Where applicable, DMSO or 5-aza-Cytidine (50µm) was added to fresh culture medium 18h before the 1st or 6th KCl-treatment. Cell density was maintained throughout the experiment by splitting them into extra dishes.

For samples for RNA-Seq only, we extracted total RNA with the GenElute™ Mammalian Total RNA Miniprep Kit (#RTN350-1KT, Sigma Aldrich, USA). For both RNA-Seq and WGBS, we extracted cytoplasmic RNA for RNA-Seq and the corresponding nuclear DNA for WGBS, using our previous nucleo-cytoplasmic fractionation protocol ^40,47^. For RT-PCR of the non-RNA-Seq samples, cytoplasmic RNA was used.

### RNA-Seq and WGBS analyses

RNA-Seq analyses were performed the same as our previous procedures ^47^, except that the Illumina HiSeq4000 paired-end 100-bp sequencing was used for the total RNA of non-treated (NT), 1^st^ KCl, or 6^th^ KCl samples, and the Illumina NovaSeq 6000 S2 paired-end 100-bp sequencing was used for the cytoplasmic RNA of the 6^th^ KCl samples with or without 24h pre-treatment by 5-azaC (50μm). Alternative exons, alternative transcription starts and alternative polyadenylation are identified by DEXSeq^47,57^; alternative splice junctions by MATS ^58^; differential gene expression by edgeR ^59^.

For WGBS analyses, approximately 1 μg of gDNA each sample was subject to bisulfite conversion for shotgun library construction (NEB Ultra II) and Illumina HiSeqX PE150 sequencing, yielding 150-bp paired-end reads. DNA quality control, library preparation, Illumina library quality control and Illumina HiSeqX PE150 sequencing were conducted at the McGill University Génome Québec Innovation Centre (Montréal, Québec, Canada). We obtained an average of 66 ± 4 million of paired-end reads. The sequence quality was verified using FastQC ^60^, with the high-quality reads mapped to the rat genome assembly Rnor_6.0.84 (GH3 samples) or mouse assembly GRCm38 (mm10, hippocampus tissue samples), using BSMAP ^17,61^. The DNA methylation status of individual cytosines of each exon was obtained by filtering the BSMAP output list with the genomic coordinates within the DEXSeq list of changed exons. Total DNA methylation level of an exon (mCpG or mCpH) was calculated by multiplying the average methylation ratio of CpG or CpH cytosines with the total number of mCpG or mCpH sites, respectively, in the sense strand of each exon.

For functional enrichment analysis and functional annotation of genes, we used the Database for Annotation, Visualization and Integrated Discovery (DAVID, developed at the U.S. National Institute of Allergy and Infectious Diseases, https://david.ncifcrf.gov) ^62^. For sequence motif analysis, the MEME was used with default parameters ^63,64^. The motifs presented are the highest scoring for tested data set.

### Semi-quantitative RT-PCR

RT-PCR was performed based on our previous procedures ^10,47^. Briefly, for reverse transcription, 300ng of cytoplasmic RNA was used in a 10ul-reaction and incubated at 45°C for 50min. For PCR, 1ul of RT product was amplified in a 12.5ul-reaction for 28-32 cycles. PCR products were resolved in 2-2.5% agarose gels containing ethidium bromide (EtBr), visualized under UV light and captured by a digital camera. Percentages of the splice variants were calculated based on band intensities quantified using ImageJ (National Institutes of Health).

Exon rank/numbers in the text/gels are based on the RGSC 6.0/rn6 assembly: *Slo1* STREX (exon), NM_031828.1; *Rps24* last exons, FN801636(upper band) & NM_031112.1(lower band); *Ehmt2* exon 9, NM_212463.1; *Dlg1* exon 20a, NM_012788.1; *Epb41l3* exon 15, NM_053927.1; Kidins220 exon 26, NM_053795.1; *Mapt* exon 6, M84156, equivalent to human *MAPT* exon 4a (NM_001123066.3, GRCh38/hg38); *Mapt* exon 7a, M84156, equivalent to human *MAPT* exon 6 (NM_001123066.3, GRCh38/hg38); *Mapt* exon 10, M84156, equivalent to human *MAPT* exon 10 (NM_001123066.3, GRCh38/hg38); *Nrg1* exon 12, NM_001271128.1; *Phldb1* exon 10, X74226. *Baiap2* exon 3, CK840478, equivalent to the mouse *Baiap2* exon 14 (ENSMUST00000106233.1, GRCm38/mm10, Figure 3).

### Splicing reporter assay

Three complementary pairs of oligo inserts are listed below with different exon methylation capacities (CpG sites underlined) were synthesized using the *mBaiap2* exon 14 as a template: (CpG)_x3_, (CpG)_x7_, and (GC)_x7m_, containing 3, 7 and 3 CpG sites, respectively, harboring the 48bp exon (uppercases) and partial flanking introns (lowercases) with ApaI (5’-GGGCC↓C-3’) and BglII (5’-A↓GATCT-3’) restriction sites at the 5’ and 3’ ends, respectively. Complementary single-stranded oligos were denatured at 95°C, 5min and annealed at 60°C 30min before restriction digestion and ligation into the vector of the splicing reporter.

(CpG)_x3_:

5’-gcgggccctgaccttgtgtttccttacagCGCGGATGTCGAAGTGGCCAGATTTTGAGCTGCCCCTG ACTAGAGTTAgtaagttgagatctatgc-3’

(CpG)_x7_:

5’-gcgggccctgaccttgtgtttccttacagCGCGGATGTCGAAGTGGCGCGATTTTGAGCTGCCCGCGACTAGAGTTAgtaagttgagatctatgc-3’

(CpG)_x7m:_

5’-gcgggccctgaccttgtgtttccttacagCGCGGATGTCGAAGTGGCCAGATTTTGAGCTGGGCCTGACTAGAGTTAgtaagttgagatctatgc-3’

The splicing reporter assay for the unmethylated and methylated dsDNA inserts was based on a reported procedure for transcription assay ^65^. Briefly, 72ug of each insert dsDNA fragment was digested by ApaI (#ER1415, Thermo Fisher Scientific, US) and BglII (#ER0082, Thermo Fisher Scientific, US), fractionated in 1% agarose gel, excised and purified using the QIAquick Gel Extraction Kit (#28706, QIAGEN, Germany). Half of the DNA was methylated *in vitro* with the CpG methyltransferase M.SssI (M0226L, New England Biolabs, USA) using S-adenosylmethionine (SAM) as a methyl group donor, and the methylation efficiency verified by the CpG methylation-sensitive restriction enzyme BstUI (restriction site: CpG↓CpG, #R0518S, New England Biolabs, USA). The insert DNA fragments with or without methylation were ligated with the same double-digested splicing reporter vector DUP175^12^, by T4 DNA ligase (Cat. # 15224-041, Invitrogen, US) at 14°C-16°C overnight. The ligated splicing reporters containing the methylated or unmethylated exons were concentrated using the QIAquick Gel Extraction Kit and co-transfected directly with the Flag-CaMKIV-dCT (CaMKIV) or –dCT-K75E (CaMKIVm) expression plasmid ^12,66,67^, into HEK293T cells using LipoFectamine 3000 (#L3000008, Invitrogen, US) and incubated overnight (16-18h) before RNA extraction.

### Lentivirus transduction and gene overexpression

The lentiviruses were purchased from GeneChem. The target sequence of MeCP2 shRNA is 5’-CAGCATCTGCAAAGAGGAGAA-3’, hnRNP L shRNA is 5’-GCTATGGTGGAGTTTGATTCT-3’, and the non-targeting control is 5’-TTCTCCGAACGTGTCACGT-3’. These sequences were cloned into the lentiviral vector GV493 containing sequences of hU6-MCS-CBh-gcGFP-IRES-puromycin. GH_3_ cells were transduced with supernatants containing virus carrying the shMeCP2 or shhnRNP L construct using the Lipofectamine 3000 reagent (Invitrogen). After 48 h, the transduction effect was verified with fluorescent microscopy, and the infected cells were then selected with 10 mg/mL puromycin for 75 days.

For MeCP2 and hnRNP L co-expression, plasmid GV366 (CMV-MCS-HA-SV40-Neomycin) was used to clone full length of hnRNP L (NM_001134760) using XhoI and BamHI restriction enzymes. The plasmid GV657 (CMV-MCS-3flag-polyA-EF1A-zsGreen-sv40-puromycin) was used to clone full length MeCP2 (NM_022673) using BamHI and KpnI restriction enzymes. The control plasmid is CON237. All constructs generated were confirmed by sequencing. HEK293T cells were transiently transduced with plasmid constructs for the overexpression of MeCP2-Flag and hnRNP L-HA using the Lipofectamine 3000 reagent (Invitrogen).

### Western blotting

Total or nuclei proteins obtained from cell lysates were subjected to SDS polyacrylamide gels, the proteins were separated by electrophoresis, and blotted onto PVDF membranes (Millipore, IPVH00010). After blocking with 5% skimmed milk, the membrane was incubated overnight with the primary antibodies at 4 °C, and then followed by horseradish peroxidase-conjugated secondary antibody and detected by chemoluminescence using ECL reagent (BIO-RAD, 1705061). The primary antibodies included anti-MeCP2 (CST, 3456), anti-hnRNP L (Santa Cruz, sc-32317), anti-METTL3 (Abcam, ab195352), anti-hnRNP F/H (Santa Cruz, sc-32310) and Anti-GAPDH (Abcam, ab181602).

### Nuclei extraction and Co-immunoprecipitation

HEK293T cells were washed twice in ice-cold PBS and resuspended in NP-40 buffer (75 mM NaCl, 0.325% NP-40, 1 mM EDTA, 10 mM Tris-Cl; pH 7.5) with Halt^TM^ protease and phosphatase inhibitor cocktail (Thermo Scientific, 78441). The suspension was gently pipetted until no visible pellet, incubated on ice for 30min, and centrifuged at 2400×g for 10 min at 4°C. The resulting pellet was washed twice with ice-cold PBS, resuspended in lysis buffer (25 mM Tris, 150 mM NaCl, 1% NP-40, 1.5mM MgCl_2_; pH 8.0) and sonicated with a regimen of ten 1-second pulses at 4°C, and treated with Benzonase (Merck Millipore, 70746) and DNase I (Thermo Scientific, EN0521) at room temperature for a 1h.

Co-immunoprecipitation for the MeCP2-Flag and hnRNP L-HA interaction was performed using a Pierce™ Co-IP Kit (Thermo Scientific, 26149) following manufacturer’s protocol. Briefly, anti-FLAG (Sigma, F1804) or anti-HA (CST, 3724) antibodies were immobilized onto the AminoLink Plus Coupling Resin. The nuclear lysate was pre-cleared with control agarose beads before the antibody-resin addition and overnight incubation at 4°C. The resin-protein complex was then washed twice in IP Wash Buffer (25mM Tris, 150mM NaCl, 1mM EDTA, 1% NP-40, 5% glycerol; pH 7.4), followed by a final wash in 1× Conditioning Buffer (pH7.2), then incubated with 40uL of Elution Buffer (pH 2.8) at room temperature for 5 minutes. The resulting elute was mixed with 10µL of loading buffer (300mM Tris•HCl, 5% SDS, 50% glycerol, pH 6.8) and heated at 100°C for 5min for western blot analysis.

### Genome/transcriptome analysis of the datasets from wild type or Rett syndrome mice

We analyzed the DNA methylation or alternative splicing of the hippocampal tissue samples of wild type or Rett syndrome mice using the raw reads from two published datasets ^19,55^. Briefly, we analyzed the raw reads of RNA-Seq sequences from the total RNA of the hippocampi of male littermate mice (wild type and Mecp2 null mice) at 7-weeks of age upon KA treatment by intraperitoneal injection ^19^, and the raw reads of WGBS from gDNA of the hippocampal dentate gyri of 8-10 week-old male mice (C57BL/6, same as the Mecp2-null background) ^55^. The reads were quality-controlled by FASTQC, trimmed and mapped to the mouse assembly GRCm38 (mm10) for DEXSeq or BSMAP analyses.

### RNA samples from Rett syndrome patients

Usage of the patient samples in the study was under the approval of the Health Research Ethics Board of the University of Manitoba. Total RNA samples were isolated from human hippocampus and cerebellar tissues using TRIzole (Life Technologies), as we reported ^68,69^. Briefly, 0.5 mL of TRIzol was added to the frozen brain powders of about 50 mg in each tube, then homogenized and incubated for 5 min at room temperature. We then added 0.1 mL chloroform, incubated it for another 3 min, and centrifuged for 15 min (12,000xg, 4°C). We collected the aqueous phase and added 5 µg RNase-free glycogen and 0.25 mL isopropanol, incubated it for 10 min at room temperature, and centrifuged for 10 min (12,000xg, 4°C). We washed the pellet with 0.5 mL of 75% ethanol and centrifuged for 5 min, 12,000xg, 4°C. RNA pellets were air-dried and re-suspended in 30µl of RNase-free water, quantified by NanoDrop 2000 micro-volume spectrophotometer, and stored at −80°C before RT-PCR. The Rett Syndrome human brain tissues used in this study are:

| NIH# | RTT | Age (years) |  | Sex | PMD |
| --- | --- | --- | --- | --- | --- |
| * | T158M | 13 | F | < 6 h |  |
| * | A201V | 19 | F | 24 h |  |
| 4516 | R255X | 21 | F | 9 h |  |
| 1815 | IVS3-2A>G | 18 | F | 5 h |  |
| 5723 | Not Available | 22 | F | 6 h |  |

\*Donated brain tissues to the Rastegar lab for research<sup>68,69</sup>.

*Donated brain tissues to the Rastegar lab for research ^68,69^.

### Statistical tests

We used two-tailed Student’s t-test, except for the built-in tests in DAVID (modified Fisher’s exact test). The DEXSeq uses Fisher’s test ^57^.

### Data availability

The raw reads of RNA-Seq of RNA, WGBS of DNA, extracted from the rat pituitary GH_3_ cells with differential treatments are available at Sequence Read Archive (SRA) database: https://www.ncbi.nlm.nih.gov/bioproject/PRJNA701032.

## Acknowledgments

We thank the University of Maryland Brain and Tissue Bank, a Brain and Tissue Repository of the NIH Biobank (at NIH NeuroBioBank Program: neurobiobank.nih.gov), and the patient families who donated brain tissues for research, and Frederick Robidoux and Sylvie Laboissiere of the McGill University Genome Quebec Innovation Centre for their excellent service. We thank Dr. Chunyu Liu for helpful comments on the manuscript. The work has been supported by a Manitoba Research Chair fund, a CIHR (Canadian Institutes of Health Research) operating grant FRN_106608, and in part by NSERC (Natural Sciences & Engineering Research Council of Canada) discovery grants (RGPIN-2016-06004 and -2022-05023) to JX, and by University of Manitoba Rady Innovation Funds to MR and JX, and NSERC-DG grant (2016-06035) to MR. LL, UD and SO by UMGF scholarships, and SO by a Vanier Graduate Scholarship.

## Supplementary Figure Legends

**S_Figure 1.**
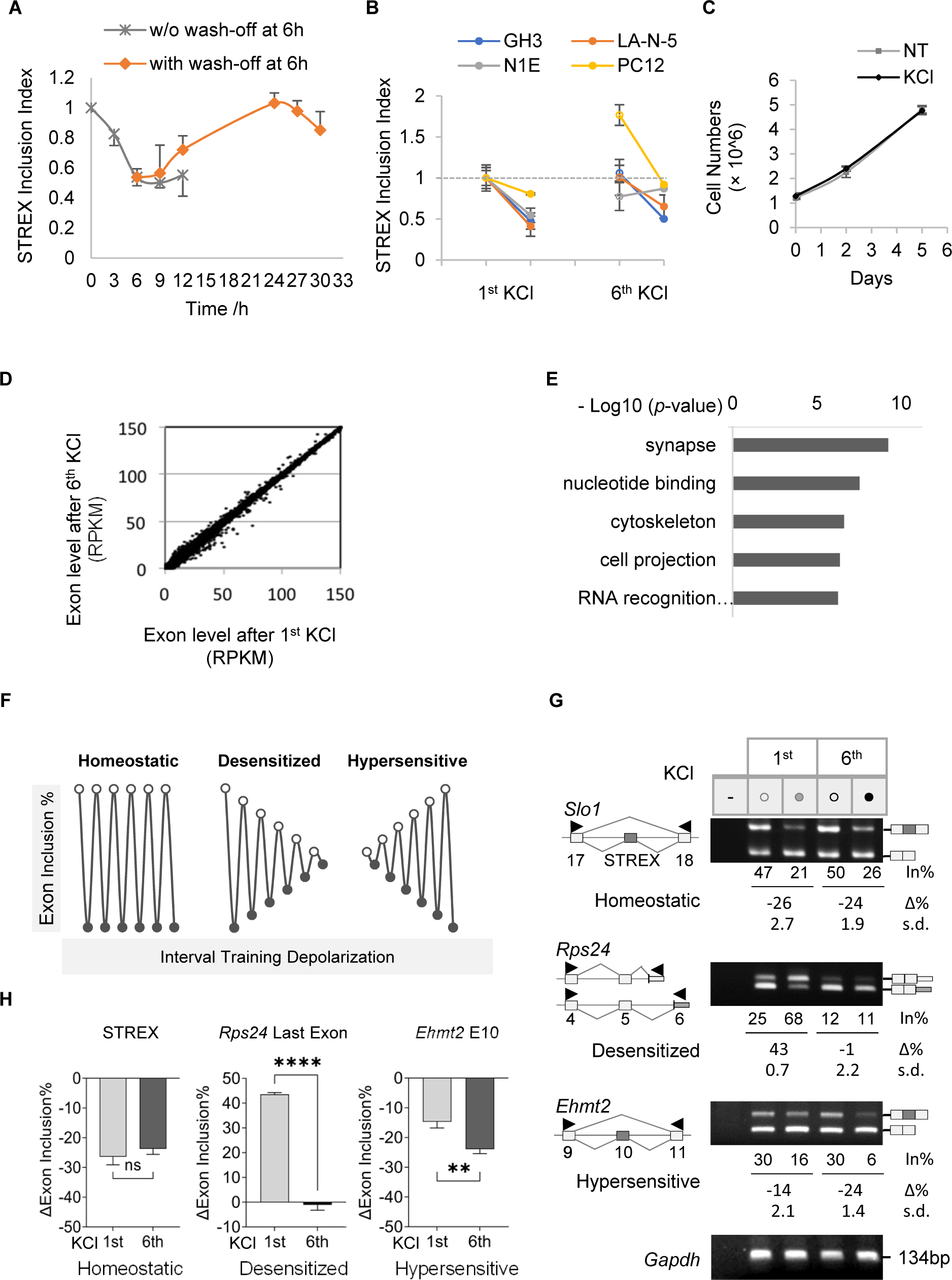
Adaptive splicing upon ITD KCl treatments: pre-tested STREX, different cell lines, exome analysis and examples of RT-PCR validations. **A.** Pre-test for the time course using STREX splicing response to single KCl (50mM) treatment (gray) or with KCl wash-off at 6h (orange) in GH_3_ cells. The STREX inclusion level before the treatment is indexed as 1. **B.** Cell-dependent adaptive splicing of STREX upon ITD (mean ± s.d., n = 3). **C**. GH_3_ growth curves throughout the treatments. Note that the STREX showed homeostatic splicing response to depolarization in GH_3_ and LA-N-5 cells here but adaptive splicing in the PC12 and N1E cells, with basal levels of 50%, 45%, 11% and 30% in the NT samples of the respective cell lines. NT: untreated. The changes in STREX inclusion levels induced by both the 1st and 6th KCl treatments were found to be statistically significant across all examined cell lines (p < 0.05). The only exception was the lack of significant change in STREX inclusion level in N1E cells following the 6th KCl treatment. **D**. Scatter plot of the exon inclusion level (normalized average reads, n = 3) between the 1^st^ and 6^th^ KCl-treated samples by DEXSeq analysis. RPKM, reads per kilobase per million. **E**. DAVID functional clusters of 1,204 genes whose exons changed between the 6^th^ and 1^st^ KCl-treated samples based on DEXSeq and MATS analysis. **F**. Schematic diagram of ideal changes according to the three primary splicing response patterns observed, with KCl-downregulated exons as examples. **G**. Representative agarose gels of RT-PCR-validated exons of the three primary splicing response patterns. Boxes in grey: alternative; white: constitutive exons; arrowhead: primers; In: inclusion of alternative exon; Δ: net change of percent exon inclusion by KCl treatment; -: PCR negative control; *Gapdh*: RNA loading control. **H**. Bar graphs of the net percent changes of the exon inclusion upon single (1^st^) or ITD (6^th^) KCl treatment (mean ± s.d., n ≥ 3). Open circles: before, and filled circles: after KCl treatment. ns, not significant; **: p < 0.01; ****: p < 0.0001.

**S_Figure 2.**
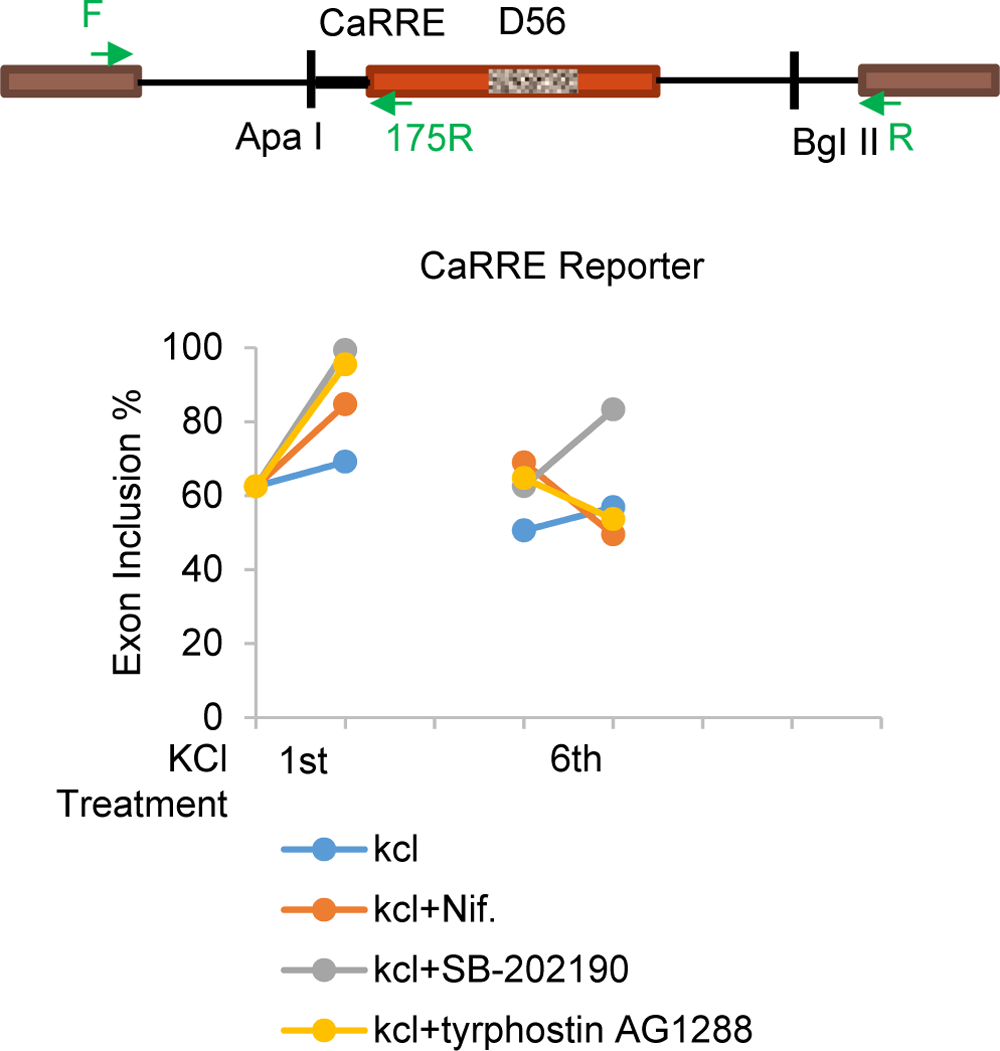
Effect of signaling pathway inhibitors on the adaptive splicing of a reporter exon. Scatter plot of tyrphostin AG 1288, nifedipine and SB202109 treatment on the depolarization effect of the 6^th^ KCl treatment on the reporter exon in N1E cells. In this assay, SB-202190 (p38 MAPK pathway inhibitor), Tyrphostin AG 1288 (Tyrosine kinases pathway inhibitor) and nifedipine (Ca2+/CaMKIV pathway inhibitor) were tested. One day after transfection of the reporter plasmid DUP175-CaRRE-D56 (CaRRE: a CaMK IV-responsive RNA element, 53-nucleotide intron + 1-nucleotide STREX exon), cells were treated with 50mM KCl and respective inhibitors (10μM each) for 6h, then washed and added back with fresh complete growth medium till the next treatment 18h later as in Fig. 1A. The assays were carried out in duplicates. Arrows: primers. The three inhibitors differentially affected both the 1^st^ and 6^th^ KCl effects on the reporter exon. Particularly, the nifedipine effect is consistent with a role of L-type calcium channels and CaMKIV as in our previous report (Xie, et al., *RNA*, 2005 Dec 11(12):1825-1834).

**S_Figure 3.**
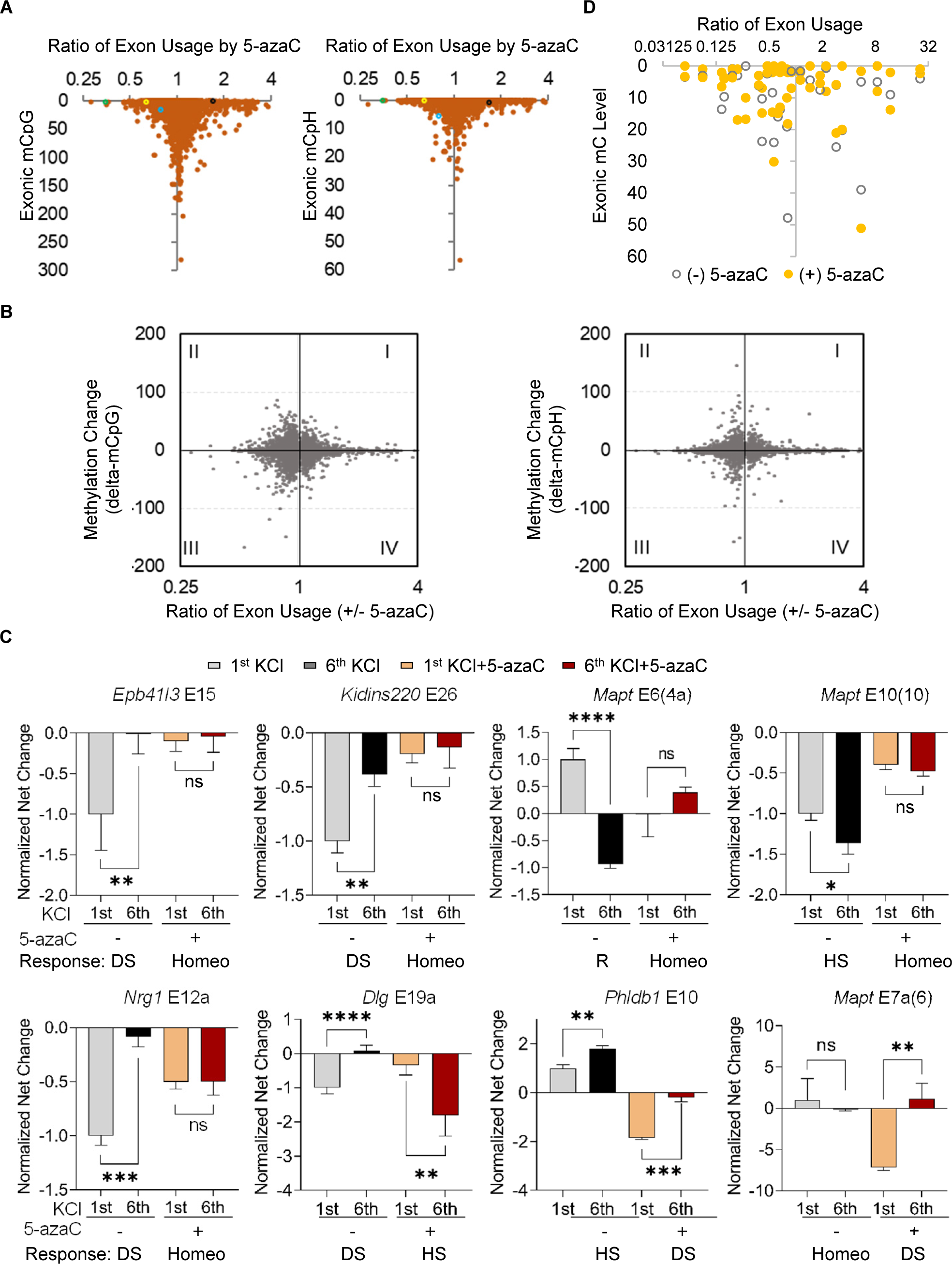
Exonic DNA methylation (EDM) levels versus splicing changes and other examples of the adaptive or homeostatic splicing of synaptic exons disrupted by 5-azaC. **A-B.** Scatter plots of the genome-wide EDM levels (A) or their index of net changes (B, per kilobases of exon DNA) and the transcriptome-wide fold changes (FC, ratio of +/− 5-azaC) of exon usage induced by 5-azaC in the 6^th^ KCl-treated GH3 cells. n = 22,361 (mCpG) and 28,077 (mCpH) exons. The total methylation level of exon DNA was calculated by the average methylation ratio (0-1) of CpG or CpH multiplied by its total number of CpG or CpH sites. In (A), examples of studied exons are highlighted: Mapt exon 6 with a yellow circle, Epb41l3 exon 15 with a blue circle, Mapt exon 10 with a green circle, and Mapt exon 7a with a black circle. **C**. Bar graphs of net percent changes of exon inclusion upon single (1st) or ITD (6th) KCl treatment (mean ± s.d., n ≥ 3) with or without 5-azaC. In brackets are the corresponding human exon rank/numbers. Homeo: homeostatic; DS: desensitized; HS: hypersensitive; R: reversed. ns: not significant, *: p < 0.05, **: p < 0.01, ***: p < 0.001. The exon numbers are based on reference transcripts in the UCSC Genome Rat Jul. 2014 (RGSC 6.0/rn6) Assembly: Epb41l3 exon 15, NM_053927.1; Kidins220 exon 26, NM_053795.1; Mapt exon 6, M84156, equivalent to human MAPT exon 4a (NM_001123066.3, GRCh38/hg38); Mapt exon 10, M84156, equivalent to human MAPT exon 10 (NM_001123066.3, GRCh38/hg38); Nrg1 exon 12, NM_001271128.1; Dlg1 exon 20a, NM_012788.1, between exons 20 and 21; Phldb1 exon 10, X74226; Mapt exon 7a, M84156, between exons 7 and 8, equivalent to human MAPT exon 6 (NM_001123066.3, GRCh38/hg38). **D**. Scatter plot of the exon mC (mCpG or mCpH) levels and corresponding ratios of exon usage (log2) by RT-PCR results, with/without 5-azaC in the 6th KCl-treated GH3 samples (n = 31 exons in total).

**S_Figure 4.**
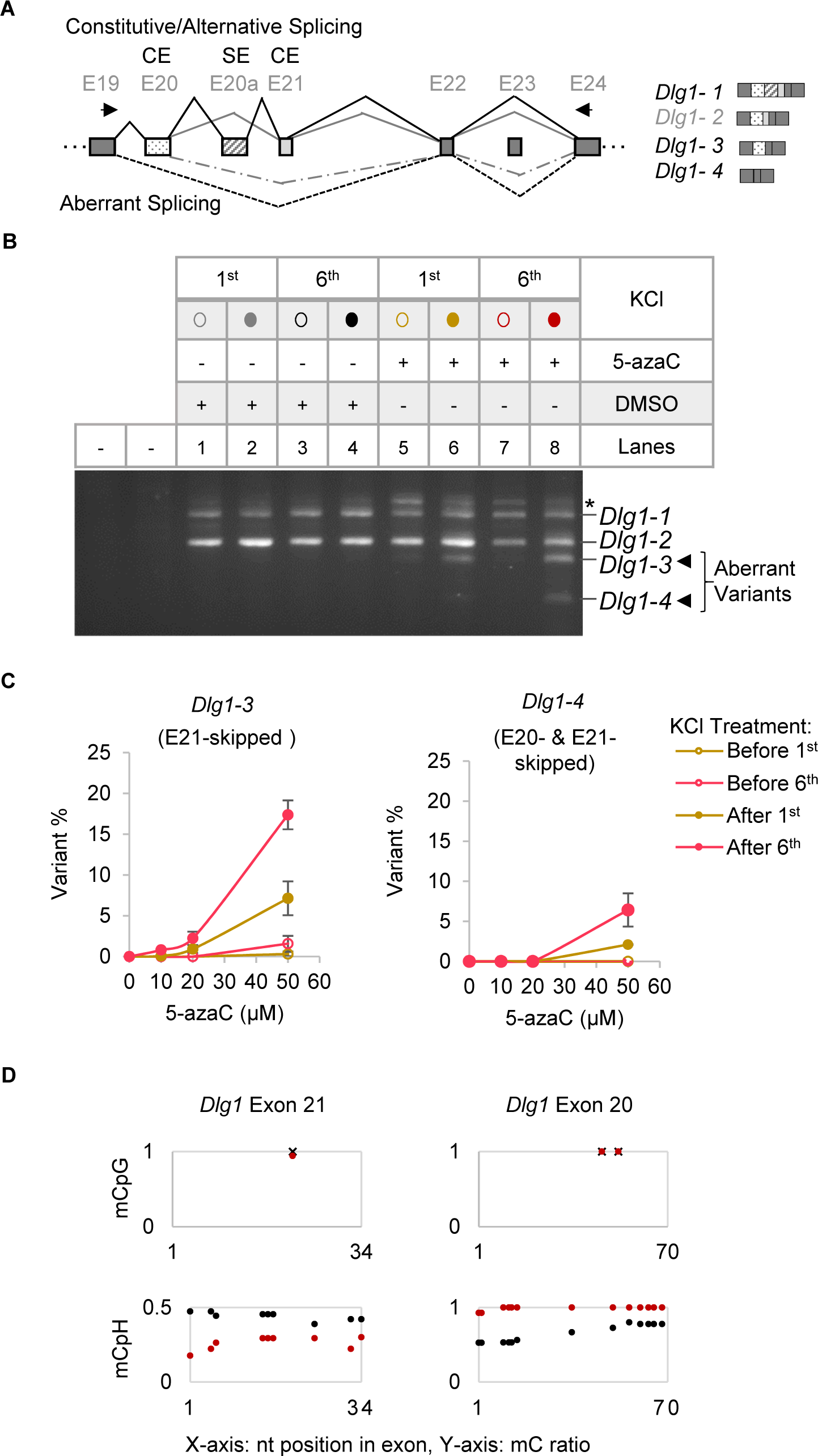
An example of the aberrant splicing of constitutive exons upon disruption of EDM by 5-azaC and its aggravation by repeated depolarization in GH_3_ cells. **A.** Diagram of the *Dlg1* splicing patterns observed after 5-zazC and ITD of GH_3_ cells. *Dlg1* exon numbering is based on the reference sequence NM_012788.1 of the RGSC 6.0/rn6 Assembly. *Dlg1* exon 20a is an alternative exon, while exons 19, 20, 21, 22, and 24 are constitutive exons in untreated GH_3_ cells. **B.** Agarose gels of RT-PCR products upon repeated KCl treatments with/without 5-azaC (50μM). Asterisk: Heteroduplex of *Dlg1-1* and *Dlg1-2*. Products were confirmed by Sanger sequencing. Circle or hollow triangle: before KCl treatment; dots or solid triangle: after KCl treatment. **C.** Dose-dependent effects on the aberrant splicing of *Dlg1* constitutive exons 21 (*Dlg1*-3 variant) and 20 (*Dlg1*-4 variant) induced by increasing concentrations of 5-azaC (0, 1nM, 1µM, 10µM, 20µM or 50µM). **D.** EDM changes of the *Dlg1* constitutive exons in the 6^th^ KCl-treated cells with (black) or without (red) 5-azaC (50µM) treatment. Upper: mCpG; Lower: mCpH.

**S_Figure 5.**
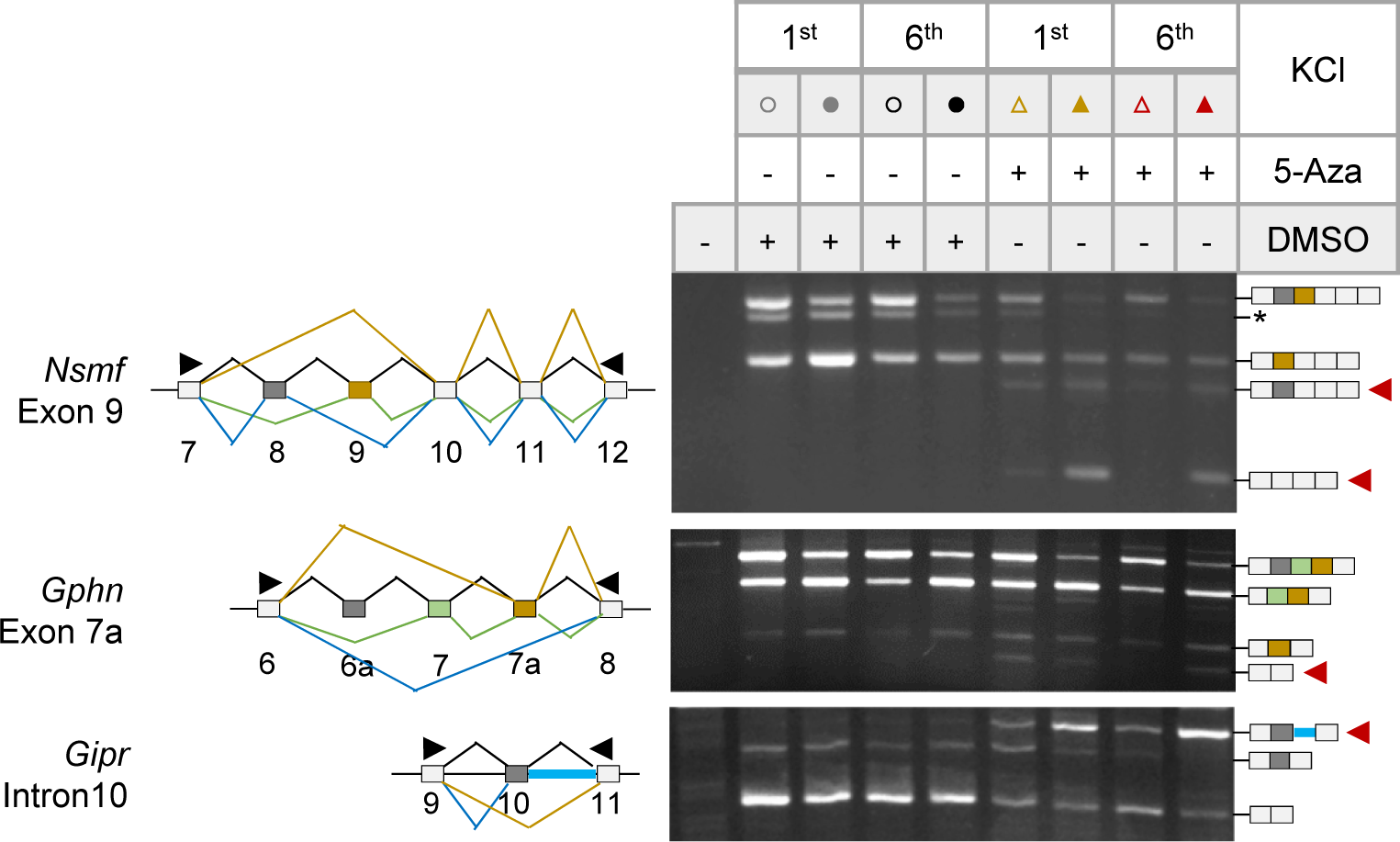
More examples of 5-azaC induced adaptively aberrant splicing of synaptic genes upon ITD. Agarose gels of RT-PCR of RNA from cells upon repeated KCl treatments with or without pre-treatment by 5-azaC (50μM) in GH_3_ cells. Open boxes: constitutive exons; Filled boxes: alternative exons; Black arrowheads: primers; Red arrowheads: PCR products with aberrant exon skipping or intron usage. Assay in triplicates. *: Product of unknown identity. The exon ranking numbers are by the reference transcripts: *Nsmf* exon 9, NM_057190.2; *Gphn* exon 7a, NM_022865.3, between exons 7 and 8; *Gipr* intron 10, NM_012714.1.

**S_Figure 6.**
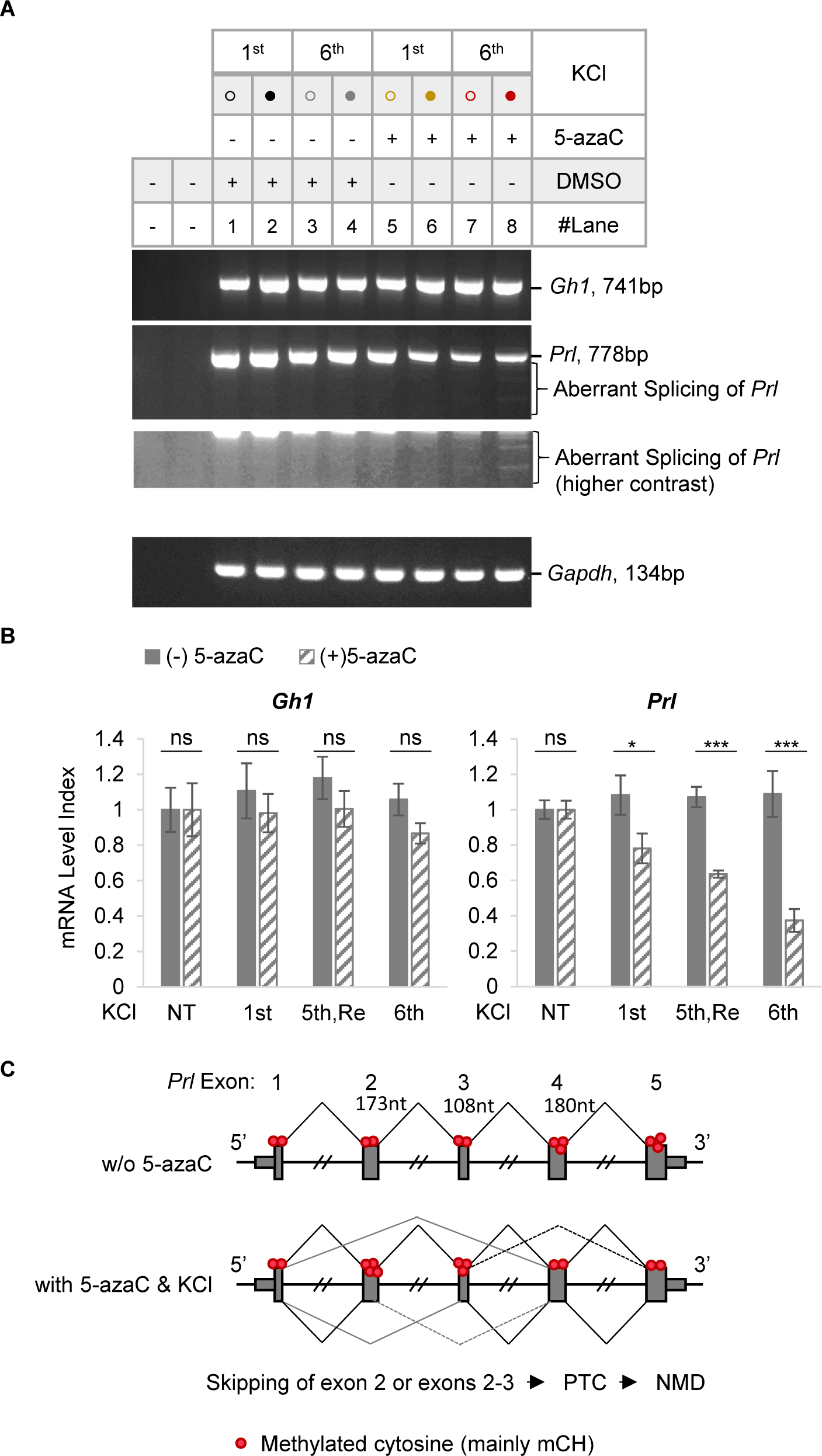
Aberrant splicing of *Prolactin* induced by 5-azaC upon ITD in GH_3_ pituitary cells, accompanied by mRNA transcript level change and EDM disruption. **A.** RT-PCR of RNA from cells upon ITD with or without 5-azaC. Without 5-azaC treatment, there are no significant change of splicing and the mRNA level of Gh1 and Prl upon either the 1^st^ or the 6^th^ KCl, suggesting that the cells have kept their endocrine identities. With 5-azaC treatment, the the *Prolactin* gene exons 2-3 were aberrantly skipped, accompanied by reduced *Prl* transcript levels and EDM changes upon ITD. *Gapdh* (Glyceraldehyde-3-Phosphate Dehydrogenase), RNA loading control. **B.** Bar graph of the normalized level of the *Prl* transcript. ns: not significant, *: *p* < 0.05; ***: *p* < 0.001. **C.** Diagram of the *Prl* variants as well as the average EDM level of each exon upon ITD with or without 5-azaC. Consistently, the EDM of these skipped exons was disrupted by 5-azaC with hypermethylation in exons 2 (from 50% to 87.5%) and 3 (from 53% to 72%), in comparison to the hypomethylation in exons 4 and 5 (from 77% to 45%). Note: the *Prl* promoter is hypermethylated (mCG and mCH, mainly mCH) from 42% to 83% upon 5-azaC treatment, suggesting transcriptional inhibition by 5-azaC via hypermethylation of its promoter. In comparison, consistent with the unchanged transcript level and stable splicing, the average methylation level of the *Gh1* promoter and EDMs weren’t significantly changed by 5-azaC.

**S_Figure 7.**
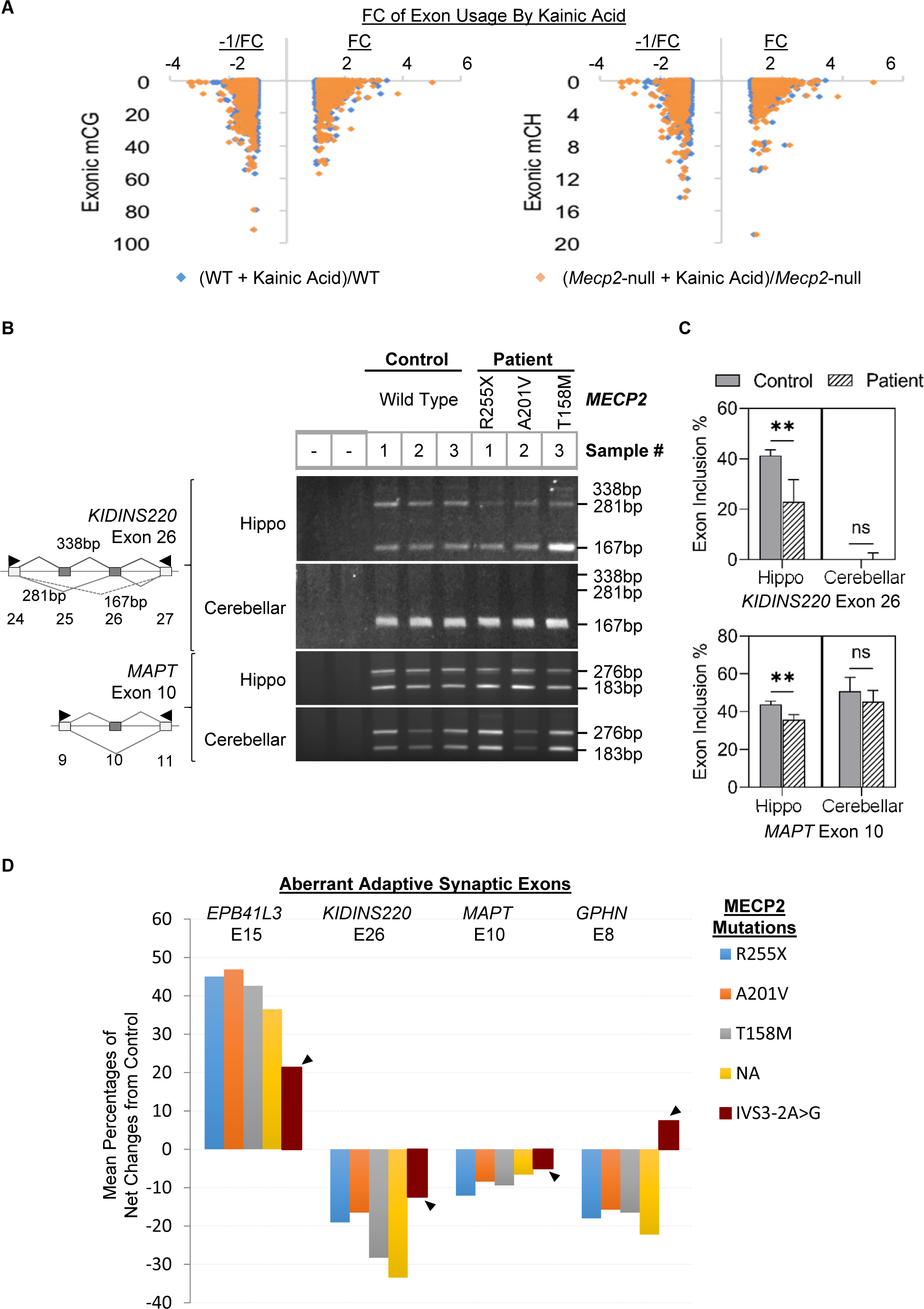
MeCP2 mutation effect on the aberrant splicing of other hippocampal genes *in vivo*. **A.** Scatter plots of the EDM mC levels of wild type mice and fold changes of corresponding exon usage with/without kainic acid treatment in the hippocampus of wild type (blue) or *Mecp2*-null (brown) mice, by analysis of the raw reads from the datasets by Osenberg or Guo respectively. The FCs less than one are displayed as −1/FC to illustrate the shift from 1 (or −1) in the *Mecp2*-null mice. Note the further shift of brown (mutant) dots overall away from the midline compared to the blues WT ones of the same group of exons. n = 1,719 exons for both mCpG and mCpH. **B-D.** Agarose gels (**B**) of RT-PCR products of splice variants, bar graphs (**C**) of their mean (± s.d.) exon inclusion levels as in Fig. 4B, and a bar graph (**D**) of the net changes of exon usage upon each *MECP2* mutation (patient) from the mean of controls. The MeCP2 IVS3-2A>G mutation consistently exhibited the smallest splicing changes compared to other mutations including R255X, A201V and T158M in different adaptive exons in the hippocampus samples of *MECP2*-mutated Rett syndrome patients, suggesting mutation-dependent differential effects on the extent of aberrant splicing. NA: mutation information not available. Arrowhead: IVS3-2A->G samples. The numbering of human exons is based on the following transcripts: *EPB41L3* exon 15, NM_012307.4, equivalent to the *Epb41l3* exon 15 (NM_053927.1); *KIDINS220* exon 26, NM_001348729.2, equivalent to the *Kidins220* exon 26 (NM_053795.1); *MAPT* exon 10, NM_001123066.3, equivalent to *Mapt* exon 10 (M84156); *GPHN* exon 8, NM_020806.4, equivalent to *Gphn* exon 6a (NM_022865.3).

**S_Figure 8.**
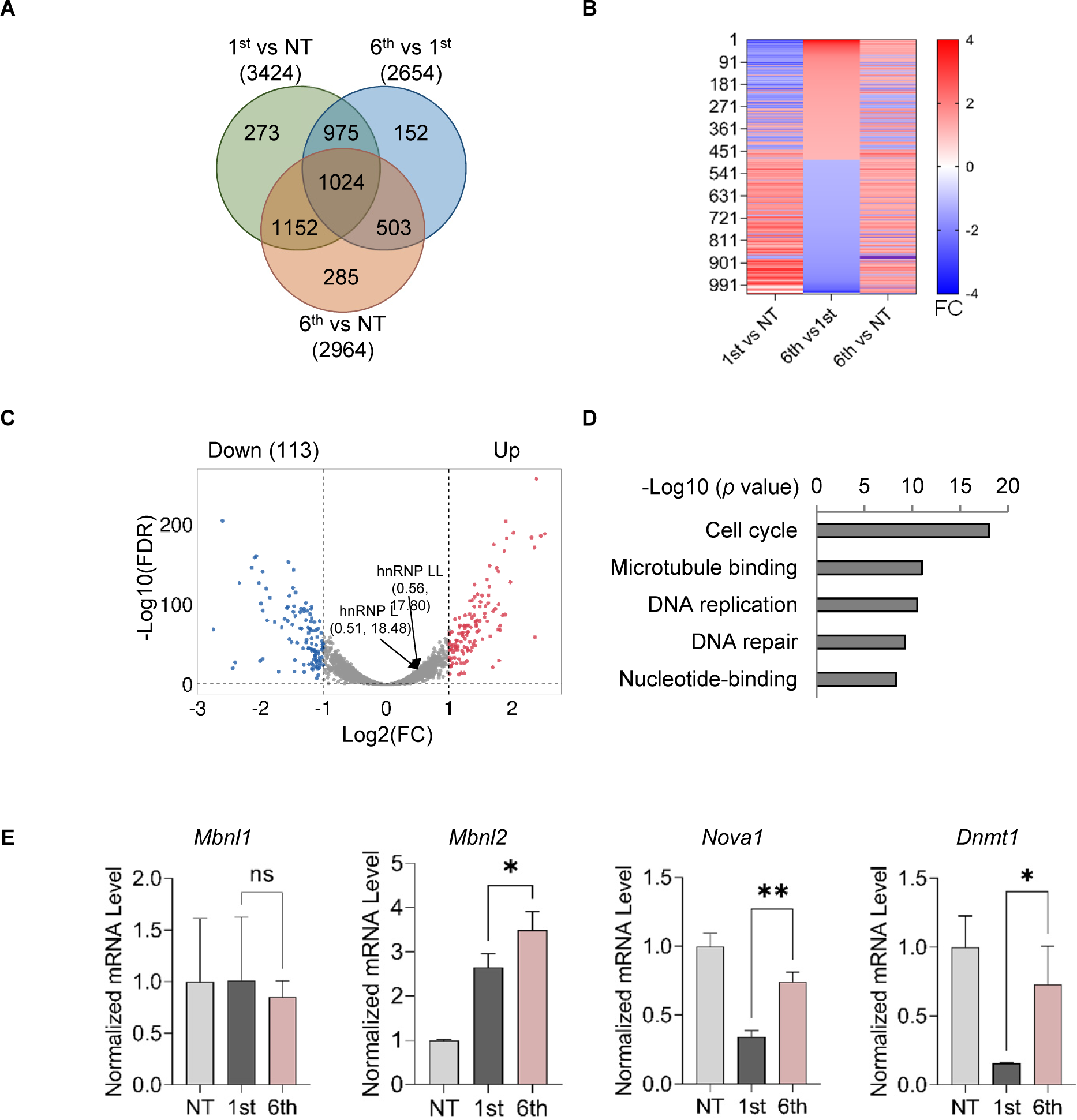
Differentially expressed genes upon ITD in GH_3_ pituitary cells. **A.** Venn diagram showing the number of differentially expressed genes upon a single (1^st^) or repeated (6^th^) KCl treatments. By filtering with average normalized counts > 50, 4,607 of 32,662 detected genes were selected for further analysis. Among them, 2,654 genes were identified with differential expression (>1.1FC) upon ITD compared to the single treatment samples, including ones (∼24.6%, 655/2654) that barely even responded to the 1^st^ treatment. **B.** Heatmap showing that 1,024 genes exhibited differential changes (>1.1FC, FDR<0.5, aveNormalized counts > 50) in transcript level between the 6^th^ and 1^st^ KCl treated samples by edgeR analysis. FC, fold change; red, upregulation; blue, downregulation. **C.** Volcano plot displaying the up-(red) and down-regulated (blue) genes (n = 238 genes, > 2FC, 1^st^ vs 6^th^ KCl treatment) upon ITD (FDR > 0.5, aveNormalized counts > 50). Black arrows indicate the position of hnRNP L and LL below the threshold, which also had no significant changes by RT-PCR in the MeCP2 knockdown cells. **D.** DAVID functional clustering analysis of the 238 differentially expressed genes, ranked by −Log10 of the *p* values. **E.** Bar graph of representative RT-PCR-validated splicing factor or DNA methylation genes with adaptive changes upon repeated KCl treatments.

**S_Table I.** Exons of synaptic genes regulated by both hnRNP L and 5-azaC, analyzed with DEXSeq and their methylation changes determined by BSMAP. In the brackets are the commonly referenced exon numbers of human *MAPT*. (Hypo: mC level reduced, Hyper: mC level increased after 5-azaC treatment, and 0 indicates no change overserved. FC: fold change.)

| Gene Name | Exon# | Gene Bank | FC by 5-azaC | FC by shL | Exonic mCpH level | Exonic mCpG level |
| --- | --- | --- | --- | --- | --- | --- |
| *Mapt* | E7a (6) | M84156 | 1.79 | -1.32 | hyper | hypo |
| *Syne1* | E90 | MK681777 | 1.56 | -1.20 | hyper | 0 |
| *Gabbr1* | E19 | NM_031028.3 | 1.27 | 1.32 | hypo | 0 |
| *Dlg1* | E20a | NM_012788.1 | -1.21 | 1.75 | hyper | 0 |
| *Cacna1d* | E46 | NM_017298.1 | -1.22 | -1.64 | hyper | 0 |
| *Epb41l3* | E15 | NM_053927.1 | -1.27 | -3.70 | hyper | 0 |
| *Mapt* | E6 (4a) | M84156 | -1.33 | 1.48 | hyper | hyper |
| *Cacna1d* | E34 | NM_017298.1 | -1.41 | -1.92 | hyper | 0 |
| *Gphn* | E6a | NM_022865.3 | -1.47 | 1.23 | hypo | 0 |
| *Clasp2* | E17 | NM_053722.2 | -1.52 | -1.18 | hypo | 0 |
| *Kifc2* | E14 | NM_198752.3 | -1.54 | -1.82 | hypo | hypo |
| *Nrg1* | E12 | NM_001271128.1 | -1.59 | 1.20 | hyper | 0 |
| *Snap91* | E29 | NM_031728.1 | -1.72 | -3.03 | NA | NA |
| *Phldb1* | E10 | X74226 | -1.89 | -3.03 | 0 | hypo |
| *Mapt* | E10 | M84156 | -2.13 | 1.21 | hypo | 0 |

**S_Table II.**
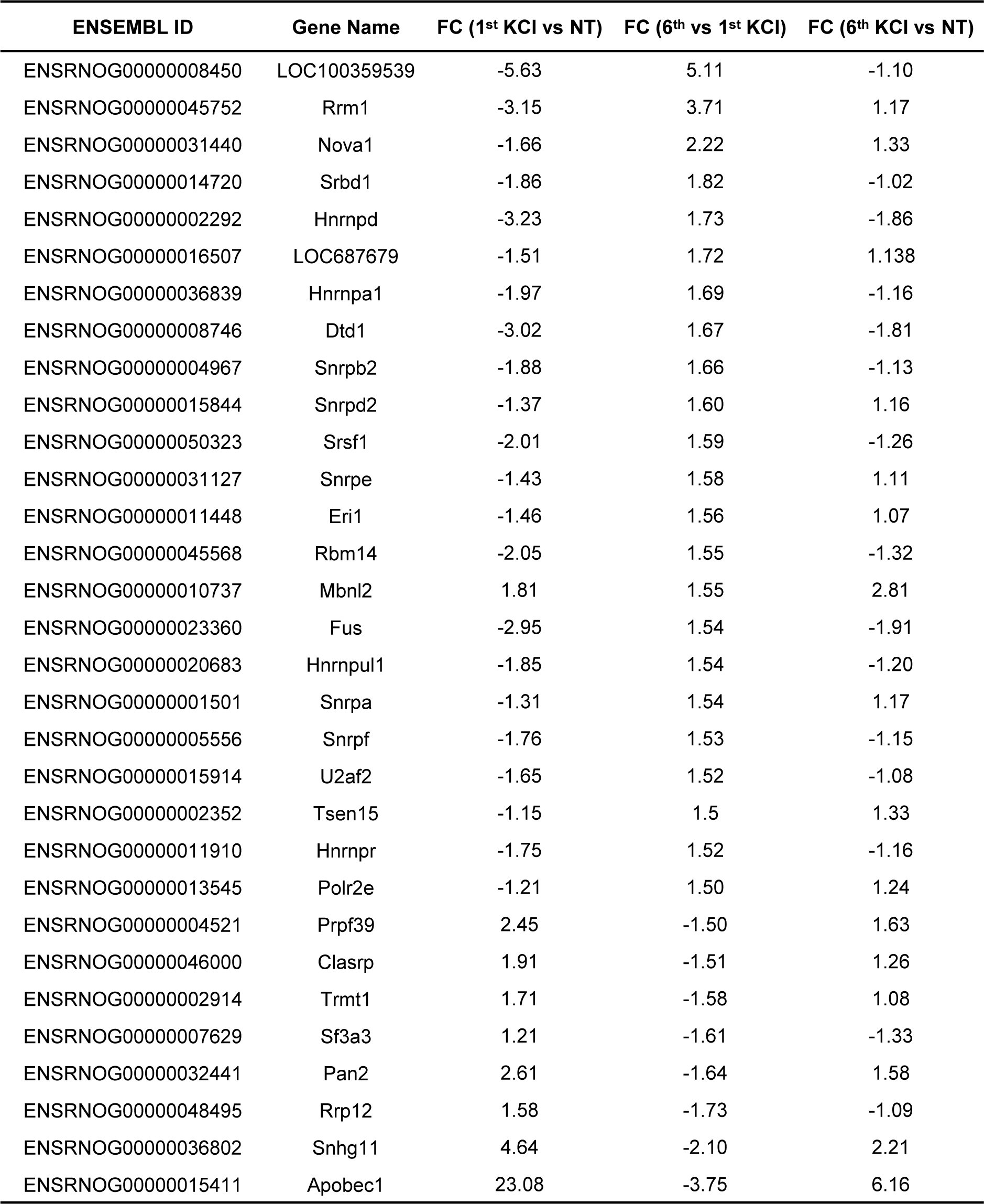
Changes in gene expression of splicing factors following repeated KCl treatment (>1.5-Fold Change between the 1^st^ and 6^th^ KCl treatment) as Identified by RNAseq.

**S_Table III.**
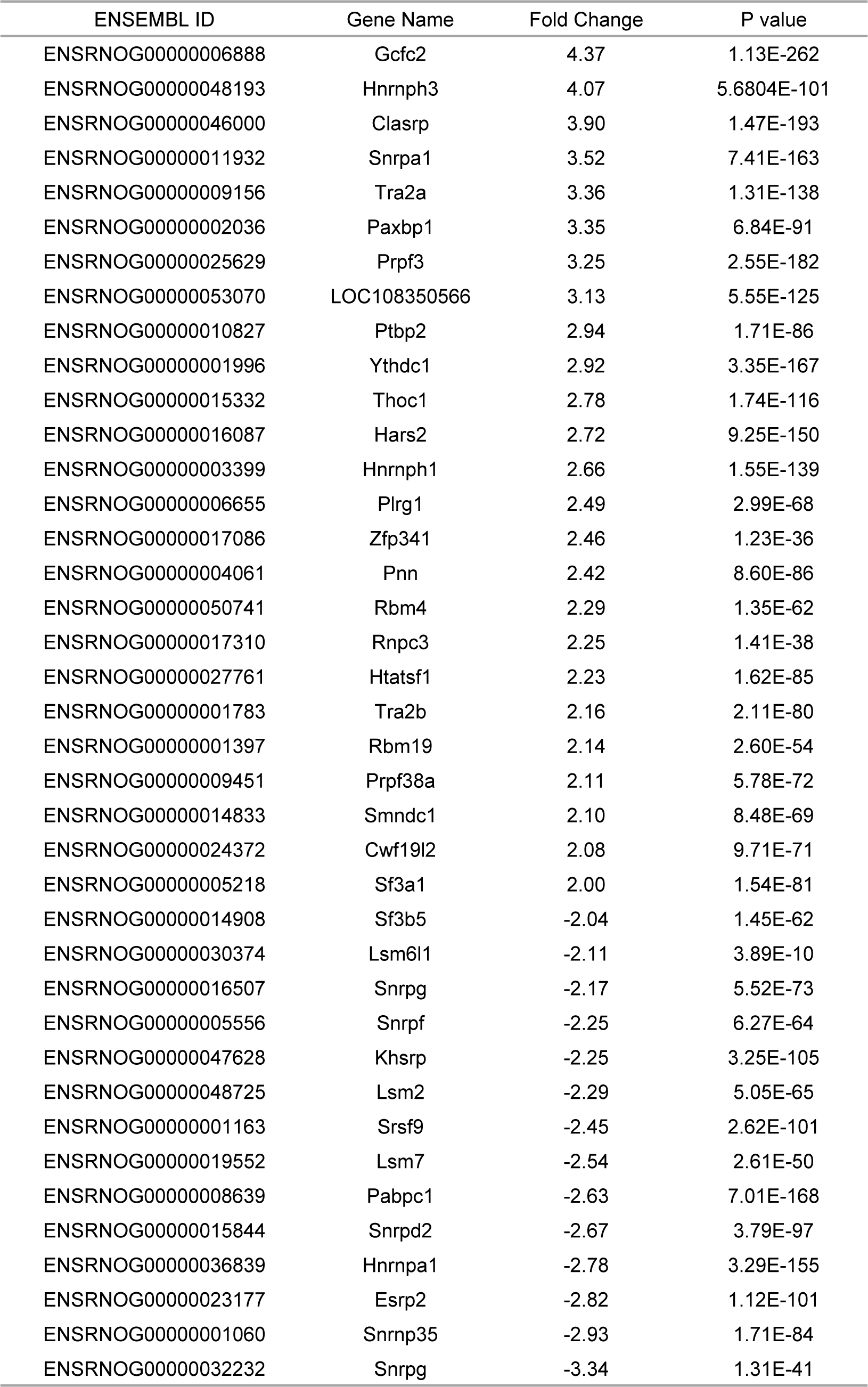
Changes in gene expression of splicing factors following 5-azaC treatment (>2-Fold Change) as Identified by RNAseq.

## Notes

### Competing Interest Statement

The authors have declared no competing interest.

